# Proton sponge or membrane fusion? – Endosomal escape of siRNA polyplexes illuminated by molecular dynamics simulations

**DOI:** 10.64898/2026.03.13.711661

**Authors:** Katharina M. Steinegger, Min Jiang, Fabian Link, Benjamin Winkeljann, Olivia M. Merkel

## Abstract

To achieve a therapeutic effect, nanoparticles delivering nucleic acids must facilitate endosomal escape (EE) of their cargo. Despite extensive research, the mechanisms that lead to an effective EE are not sufficiently understood. Herein, we utilized Molecular Dynamics (MD) simulations in All Atom (AA) and Coarse Grained (CG) resolutions to differentiate the interaction of four polymeric formulations (polyplexes) and one lipid nanoparticle (LNP) with endosomal membranes. On the one hand, the results emphasize the benefit of hydrophobic residues in the nanoparticles. On the other hand, the role of anionic lipids in the biological membranes is demonstrated. Furthermore, the identified interaction patterns were successfully correlated to the *in vitro* performance of the formulations. For the first time, different EE mechanisms of polyplex formulations are visualized in simulation and therefore distinguishable from one another. Hence, this work highlights the power of MD simulations for taking a big step towards better understanding EE efficiency.

**TOC:** 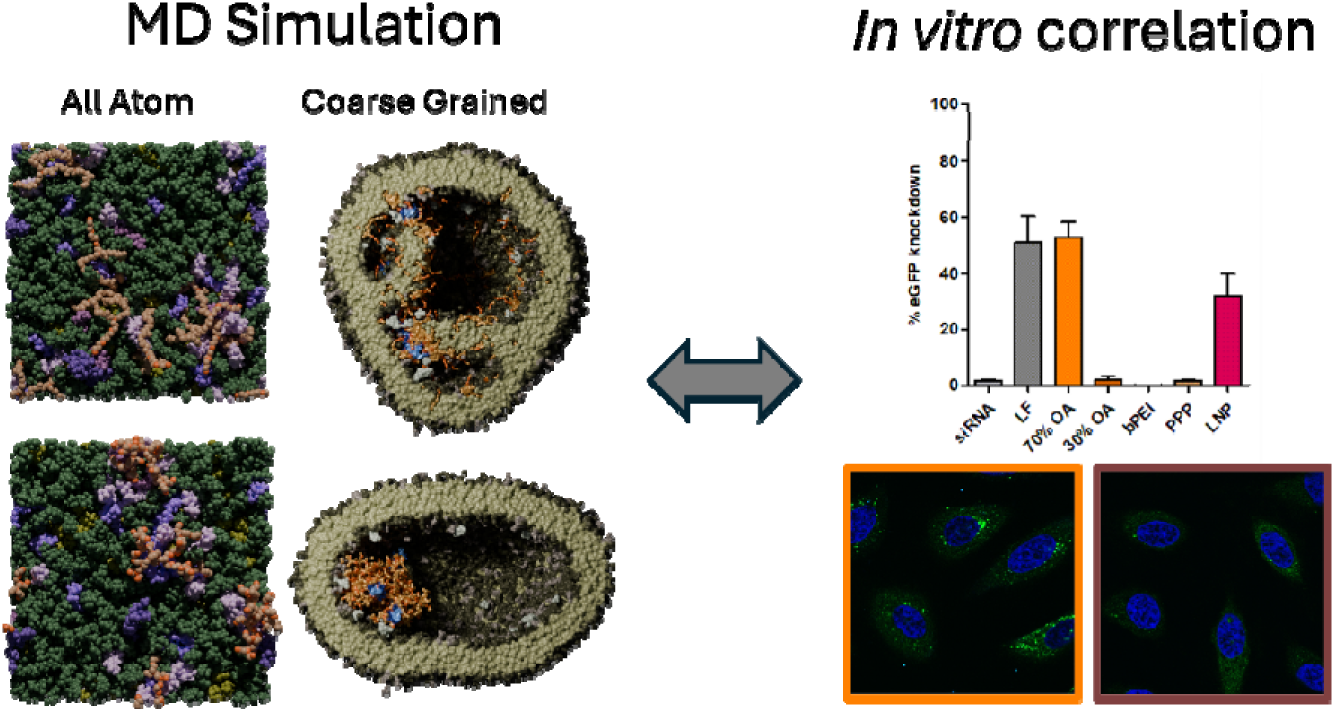

## Introduction

Short interfering ribonucleic acid (siRNA) downregulates the expression of targeted, disease-driving genes^1^ by binding to the RNA-induced Silencing Complex (RISC) in the cytoplasm. Therefore, following endocytosis, escape from the endo-lysosomal pathway is essential to achieve a therapeutic effect^2^. A broad range of strategies has been developed to formulate potent nanoparticles for nucleic acid delivery^3, 4^, including the use of polycationic polymers^5, 6^. However, many of these polymer-based polyelectrolyte complexes, commonly termed polyplexes, exhibit limited endosomal escape (EE)^7, 8^, leaving room for significant technological improvement^9, 10^.

Various mechanisms have been discussed to play a role in the EE of polymeric nanoparticles: Initially, the “proton sponge” theory, first formulated in the 1990s^11^, was among the most popular theories. It relies on the buffering capacity of endocytosed polymers, which promotes increased proton influx into the endosome during acidification. This is thought to be accompanied by the influx of neutralizing chloride ions and water^12^, resulting in osmotic swelling. Ultimately, the endosome ruptures and nucleic acid cargos could be released into the cytoplasm. A closely related hypothesis proposes that endosomal acidification increases the charge density of the polymer, leading to polyplex swelling and inducing a steric burst of the endosomal membrane^13^. These hypotheses are supported by findings that polymers with p*K*_a_ values in the physiological range (approximately 6-8) are more effective at facilitating EE^14^ and the circumstance that EE can be reduced by inhibiting endosomal acidification. However, conflicting results from live-cell imaging demand a revision of the proposed mechanisms^15^, as the data does not indicate complete lysis of endosomes after successful EE of the nucleic acid. In either case, rapid and intense disruption of endosomes and lysosomes can induce cytotoxicity due to the concurrent release of harmful vesicular contents, and is therefore considered undesirable^16^. Subsequently, attention shifts to direct polymer-membrane interactions that locally form smaller endosomal holes or pores^17, 18^. Polyplexes do not escape the endosomes intact^15^, but rather in a disintegrated state. The role of acidification could therefore be attributed to its involvement in nucleic acid cargo unpacking and polymer shedding from the particle^19^.

Concerning lipid-based nanoparticles (lipoplexes and lipid nanoparticles (LNPs)), research indicates EE to be a complex procedure including membrane fusion, phase transition in the lipid phase of the LNP, and lipid mixing between membrane and nanoparticle ^15, 20–23^.

In consequence, the lipid bilayer of the membrane is disturbed and the cargo escapes through resulting holes. LNPs can be highly effective and have successfully entered the market, for example in the form of Onpattro® (Patisiran)^24^. Still, LNPs too are limited by their EE, and only a small, often cited as single-digit percentage^25^ of the encapsulated siRNA molecules reaches the cytoplasm^26^.

Even though great effort has been put into understanding EE mechanisms, the process is still not understood to a level that allows specific fine tuning of EE performance of a formulation. As Molecular Dynamics (MD) simulations help to understand underlying mechanisms in complex formulations or biological interactions on a molecular level^20, 27^, they possess great potential to overcome the gap in understanding EE. All Atom (AA) MD simulations, showing the molecules in single atom resolution, provide detailed insights on e.g., binding mechanisms on a small scale^28, 29^. In contrast, Coarse Grained (CG) MD simulations work with a decreased resolution, as they summarize groups of atoms in predefined beads^30^. CG MD enables larger simulations up to ∼ 100 nm side length of the simulation box with longer timescales in the range of multiple microseconds-enabling for example the simulation of the formation of whole nanoparticles^31, 32^. Additionally, it has previously been shown that CG MD can visualize EE mechanisms^20^.

This work compares the EE of four polyplexes and one LNP formulation through AA and CG MD simulations. The first polyplex material is 25 kDa branched PEI (bPEI), which is a commercially available polymer that has been used for siRNA delivery in research for over 20 years^11, 33^. The second polymer is a block copolymer consisting of two blocks of 5 kDa bPEI, linked by a 5 kDa polycaprolactone (PCL) chain^34^. Furthermore, two variants of an amphiphilic poly(beta)aminoester (PBAE) copolymer^9, 35^ were tested for their EE performance. They differ in their content of hydrophobic oleylamine (OA) residues, so that a more hydrophilic particle is compared to a more hydrophobic variant. Lastly, an Onpattro®-like LNP formulation^36^ was incorporated in the study to directly compare the EE of polyplexes and LNPs.

*In vitro* experiments outlined strong differences in the performance of the compared formulations. Based on the interaction patterns visualized and identified by MD, these differences can be meaningfully interpreted. Hence, this work highlights the power of MD simulations for taking a big step towards better understanding EE mechanisms.

## Results and Discussion

### Characterization of nanoparticles

Four polyplex formulations described in the literature were included in this study and based on the polymers shown in Fig. 1A. For the PBAE, either a 70% OA polymer (i.e., more hydrophobic) or a 30% OA polymer (i.e., more hydrophilic) were used. All polyplexes were formulated at N/P 10, as all four polymers have been shown to form stable particles at that ratio^9, 33, 34, 37^. The LNP was formulated with an Onpattro®-like composition, but at N/P ratio of 6.5 to align as closely as possible with the simulated model LNP^36^. For all experiments, the formulations were normalized to equal siRNA concentration. However, due to differences in polymer charge density, the total polymer mass concentration varied notably between formulations, despite equivalent N/P ratios. Specifically, relative to bPEI, the concentrations were 7.9-fold higher for 70% OA PBAE, 5.9-fold for 30% OA PBAE, and 1.6-fold for PPP.

**Figure 1.**
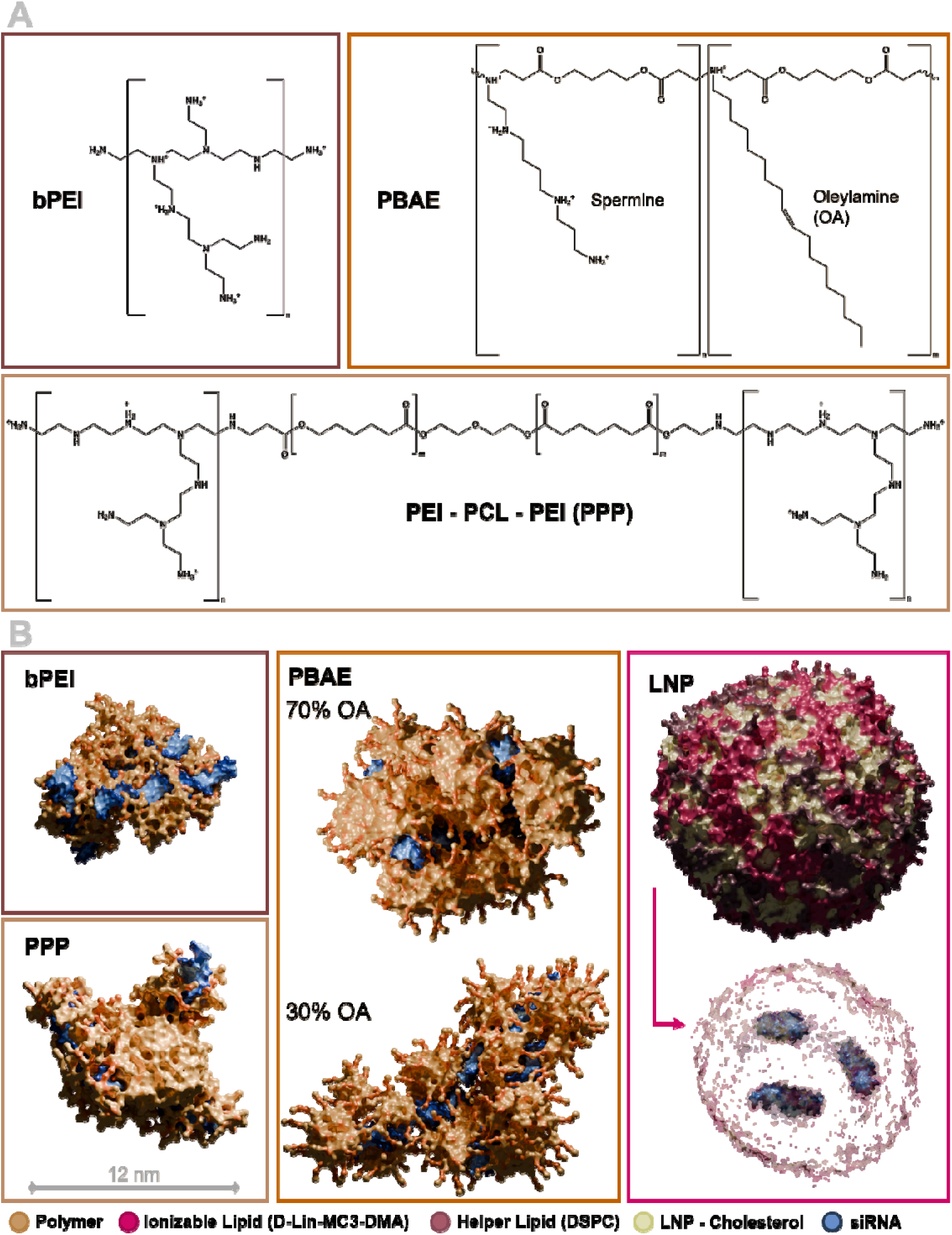
Polymer structures and model particles in CG resolution. **A.** Molecular structures of polymers used in this work; branched polyethylenimine (bPEI), poly(beta)aminoester (PBAE) with varying ratios of hydrophilic spermine and hydrophobic oleylamine (OA), and bPEI – polycaprolactone (PCL) – bPEI block copolymer (PPP). **B.** CG model particles with three siRNA molecules each at pH 7.4. The LNP is additionally shown with transparent lipids to visualize the orientation of the siRNA molecules inside; polymer in gold/orange, MC3/MC3H in pink, DSPC in purple, cholesterol in green, RNA in blue.

All five formulations formed particles with a hydrodynamic diameter (z-average) between 50 and 70 nm and a polydispersity index (PDI) below 0.3 (Fig. S1A). The ζ-potential of all polyplexes was positive, whereas the LNPs were slightly negatively charged (Fig. S1B).

In CG simulations, polyplexes were formed via self-assembly, resulting in stable nanoparticles with diameters ranging from approximately 10 to 18 nm (Fig. 1B). All polymer molecules present were associated with the respective polyplex, except for 30% OA PBAE. Here, 20% of the polymer remained separate from the polyplex at both pH 5.4 and pH 7.4, resulting in a final N/P ratio of approximately eight. This agrees with our previous work on PBAE polyplexes, where 30% OA PBAE showed unbound polymer at pH 5.4 and N/P ratios above ∼ 6^31^. For subsequent simulations of the membrane interaction, excess polymer in the 30% OA PBAE MD setup was removed. The model LNP with a diameter of ∼ 16 nm did not originate from a self assembly simulation, but was constructed based on a protocol for the setup of LNPs with hexagonal core structure in Martini 3^36^. As Polyethylenglycol (PEG) lipids tend to shed from LNPs when in contact with serum^38^ they are not expected to play a role in EE of LNPs. Therefore, no PEG lipids were incorporated into the simulated LNP model.

### Comparison of the particles *in vitro*

In HeLa cells stably expressing enhanced green fluorescent protein (eGFP), the 70% OA PBAE polyplex and the LNP showed the highest knockdown efficiencies with 30 - 60% at a dose of 20 pmol/ 6,000 cells. (Fig. 2A), whereas the other polyplex formulations achieved no knockdown at the same dose. No cytotoxicity was evident for the 30% OA PBAE and the LNP formulation, with cell viability being > 90% and lactate dehydrogenase (LDH) release being < 10% (Fig. S3) at all tested particle concentrations. The bPEI and PPP particles only caused mild cytotoxicity (< 90% cell viability) at the highest concentration. However, the 70% OA PBAE polyplexes caused notable LDH release (12-25%) and reduced cell viability (50-80%) at all tested concentrations.

**Figure 2.**
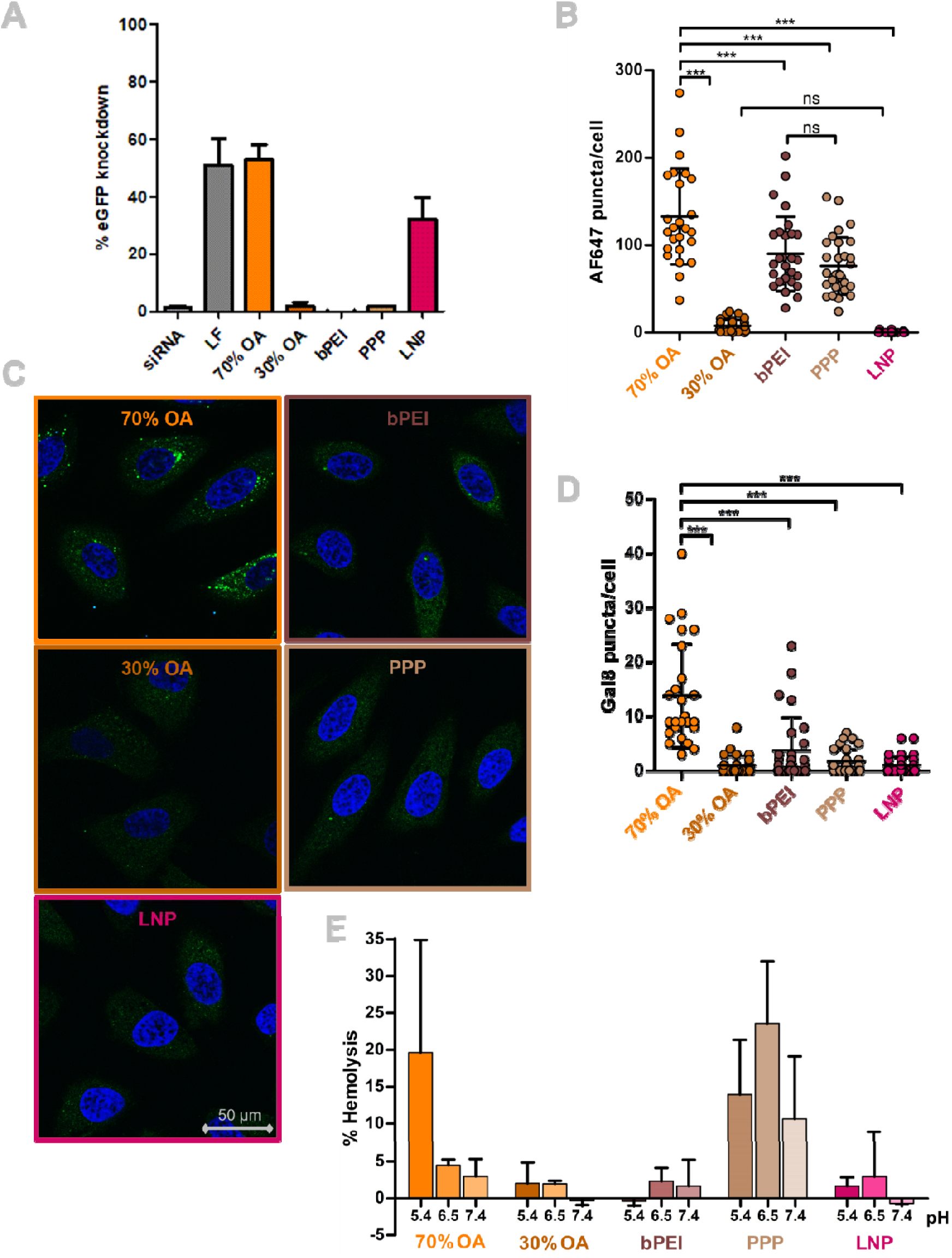
Comparison of *in-vitro* behavior. **A.** eGFP knockdown in HeLa/eGFP cells, mean ± sd, n = 2. **B.** Uptake in HeLa cells quantified as puncta of AF647 labeled siRNA per cell observed in the confocal images, mean ± sd, one-way ANOVA, ***p < 0.001, ns = nonsignificant (p > 0.05). **C.** Confocal images showing Gal8 recruitment (green puncta) in HeLa cells (blue: DAPI) 4 h after transfection. **D.** Quantification of Gal8 puncta as shown in C, mean ± sd, one-way ANOVA, ***p < 0.001, ns = nonsignificant (p > 0.05). **E.** Erythrocyte lysis of all five nanoparticle formulations relative to Triton X treatment at three different pH values (5.4, 6.5 and 7.4), mean ± sd, n = 3.

The limited knockdown efficiencies observed for the 30% OA PBAE, bPEI, and PPP polyplexes can be partially attributed to insufficient cellular uptake (Fig. 2B and Fig. S4). Among all formulations, the 70% OA PBAE polyplex consistently showed superior uptake, while 30% OA PBAE exhibited particularly low internalization, and bPEI and PPP polyplexes performed comparably to each other. The relatively weak uptake signal of the LNP in confocal microscopy was likely due to fluorescence quenching within the dense core of the nanoparticles^39^. Since the microscope settings were optimized to detect the strong fluorescence signal of AF647-labeled siRNA in polyplexes, they were suboptimal for capturing the quenched signal from the LNPs. Consequently, Fig. S4 indicates that cellular uptake of the LNPs is comparable to that of bPEI and PPP. Differences in uptake may explain the superior knockdown efficiency of the 70% OA PBAE. However, they do not fully account for the performance gap between the LNP and the bPEI or PPP polyplexes. Subsequently, EE efficiency was quantified through confocal fluorescence microscopy as puncta caused by the recruitment of mRuby-3-Galectin 8 fusion protein (Gal8) stably expressed by the cells. Gal8 is recruited to damaged endosomes when luminal glycans are exposed to the cytoplasm (Fig. 2C+D) and is therefore widely used as EE marker^40^. The 70% OA PBAE polyplexes caused significantly more Gal8 recruitment compared to all other formulations. However, these polyplexes also achieved the highest uptake by the HeLa cells. Therefore, the high knockdown efficiency of the 70% OA PBAE was likely a combination of superior uptake and strong EE. The endo-lysosomal membrane disruption caused by the EE of the 70% OA polyplexes can trigger apoptosis or uncontrolled cell death^41, 42^, consistent with the observed cytotoxicity of 70% OA PBAE. HeLa cells are characterized by relatively small endosomes, which has been suggested to favor the EE efficiency of polyplex formulations^7^. Hence, because Gal8 recruitment is cell type-dependent^43^, the same polymer may be safe and effective in other cell types^37^. The other polyplexes induced significantly lower Gal8 recruitment, suggesting that limited EE may contribute to their poor knockdown efficiency, which is in accordance with published results about PEI polyplexes^43^.

Due to the lack of cellular uptake of 30% OA PBAE, their performance in the Gal8 assay could not be directly compared to the other formulations. Rui et al.^43^ reported a negative correlation between the hydrophobicity of PBAE polyplexes and Gal8 recruitment. Notably, in their study, increased Gal8 recruitment did not translate into improved transfection efficiency. The authors suggested that high Gal8 puncta counts might result from empty polymer micelles that disrupt endosomes without delivering nucleic acid cargo. For other amphiphilic PBAEs, a positive correlation between hydrophobicity and Gal8 recruitment was observed^44^, with the most hydrophilic polymer causing the least Gal8 puncta.

Interestingly, the LNP formulation caused Gal8 recruitment as low as bPEI and PPP, even though its knockdown efficiency was higher. However, the low Gal8 recruitment by the LNPs likely originated from the EE mechanism itself: It has been shown that the ionizable lipid DLin-MC3-DMG does not induce Galectin recruitment^23^, arguably due to the formation of only small pores in the endosomal membrane. In summary, although useful within narrowly defined particle libraries^8^, Gal8 recruitment alone is not sufficient to predict the EE or knockdown efficiency across diverse nanoparticle formulations. The detection of Gal8 puncta does not indicate whether endosomal damage was accompanied by the release of nucleic acid cargo^43^, nor does it capture smaller membrane defects^23^. This could lead to favoring potentially cytotoxic particles with high Gal8 recruitment over similarly effective particles that cause only minor endosomal damage through other EE mechanisms.

Therefore, nanoparticles-membrane interactions were next assessed by measuring erythrocyte lysis (Fig. 2E). The LNPs, 30% OA PBAE and bPEI polyplexes only caused minor hemolysis, independent of the medium’s pH level. In contrast, PPP particles caused notable hemolysis of ∼ 10 – 25% at all pH levels, and 70% OA PBAE polyplexes caused increased lysis at pH 5.4 only. Hemolytic activity is generally associated with cytotoxicity^45^. However, lytic activity at acidic pH only, as demonstrated by 70% OA PBAE, is favorable for polyplexes^8^, as it indicates increased membrane interaction in the acidified endo-lysosomal compartment. Again, this emphasizes that the toxicity of the 70% OA PBAE can be referred to its excessive effect on the endosomes. As the lytic activity of the PPP polymer exceeds the cytotoxicity observed in other assays (Fig.S3), it can be related to membrane interactions in the hemolysis assay that are otherwise masked by a protein corona around the nanoparticle formed in serum^46^.

As the uptake efficiency of the formulations differed strongly, differences in EE performance could not be clearly interpreted. However, the 70% OA PBAE and the LNP formulation overall outperform the other polyplexes concerning knockdown efficiency, which will be correlated to their membrane interactions and EE mechanisms by MD below.

### Membrane interaction mechanisms in simulation

To identify the interaction mechanisms of nanoparticles with membranes after cellular uptake, AA and CG membranes mimicking different stages of the endo-lysosomal pathway were created (Table S1). The first membrane represents the (early) endosome, which is slightly negatively charged and contains glycolipids that were initially present at the outer leaflet of the plasma membrane^47^. As the milieu in the early endosome is only slightly acidic, the particle interactions with this membrane type were simulated at pH 6.5. Deemed crucial for the functionality of biological membranes is also the formation of microdomains such as lipid rafts^48^. These specified membrane regions form through the preferred interaction of cholesterol, saturated lipids and sphingolipids. To investigate the influence of lipid raft formation on EE, a simplified membrane model, composed of palmitoyloleoylphosphatidylcholine (POPC), cholesterol, and N-Palmitoyl-D-sphingomyelin (DPSM) was created. This model contained an increased amount (2%) of glycolipids (glycolip-monosialotetrahexosylgangliosides DPG1 and DPG3), which comprise one n-acetylneuraminic acid each. Hence, the lipid raft model contained all its negative charge in the glycan layer. Lipid rafts are characterized by reduced lateral diffusion^49^ in the ordered state, which was well represented by our models (both AA and CG) in comparison to the other membrane types (Fig. S2).

Two additional membrane models were constructed to simulate the late endosome/lysosome with interactions at more acidic conditions (pH 5.4). In the late endosome, the amount of cholesterol and sphingolipids was decreased, whereas the amount of negatively charged lipids was increased^47, 50^. These negatively charged lipids include bis-(monoacylglycero)-phosphate (BMGP)^51^, a lipid that is unique to late endosomal/lysosomal membranes. The late endosomal membrane contained the same number of glycolipids as the endosomal membrane used in this study. The lysosomal membrane model was structurally identical to the late endosomal model, except that glycolipids were omitted to allow assessment of their specific influence. As a result, the lysosomal membrane was the only symmetrical bilayer among all models examined. While this study aimed to reproduce the lipid composition of endo-lysosomal membranes with greater compositional diversity than previous models^20, 52^, the incorporation of membrane proteins and active cellular processes remains beyond its scope. Consequently, the presented membrane models should still be regarded as simplified representations.

The interaction of the four CG membrane models with all five particle formulations was simulated in triplicates. Initial contact of the particles with the glycosylated leaflet of the membrane was assured by a short pull applied to the particle in the first nanoseconds of simulation. After 2.5 µs, distinctive differences between the particles were detected (Fig. 3A-E). Two types of interaction with the membranes were prominent: Firstly, hydrophobic interactions of the PBAEs, the PPP polyplex and the LNP were visible as interference of polymers/LNP-lipids with the hydrophobic core of the membranes. In the case of the PBAEs (Fig. 3A+B) and the LNP (Fig. 3C), this led to mixing of membrane lipids and particle material^52^, resulting in notable amounts of particle material being shed from the particle and integrated into the membrane. Simultaneously, the 70% OA PBAE polyplex and the LNP were capable of extracting membrane lipids out of the initial membrane plane. Due to the copolymeric structure of the PBAEs, their OA tails reached into the hydrophobic membrane center, while the backbone stayed in the headgroup region and the polycationic spermines reached into the solvent layer above. However, unlike the 70% OA PBAE and the LNP, the 30% OA polymer was not capable of extracting lipids from the membrane into hydrophobic particle compartments above the membrane surface. Hence, the interaction mode of the hydrophilic PBAE appeared more comparable to the influence of bPEI and PPP, which caused only minor to no deviations of the membranes’ density profiles (Fig. 3B, D and E). In the PPP polymer, the hydrophobic compartment of the particle (the PCL chain) is clearly separated from the hydrophilic bPEI units due to the block copolymeric structure. Consequently, if in immediate contact with the membrane, the PCL residue tended to partition into the membrane core, whereas the bPEI segments remained localized at the membrane surface (Fig. 3E, Fig. S5).

**Figure 3.**
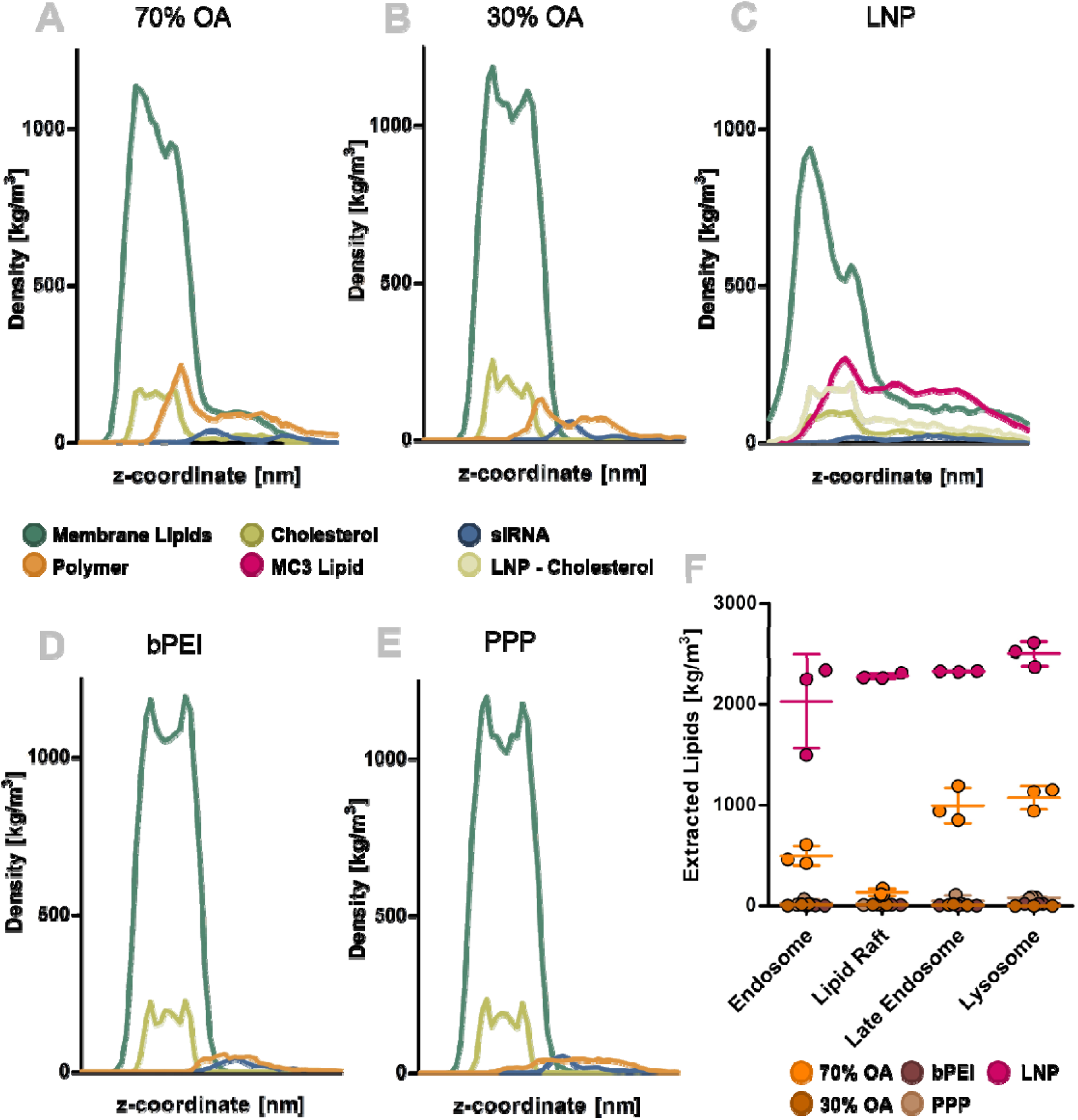
CG-MD simulations of nanoparticles interacting with planar membrane models. Density distribution along the z – coordinate of the simulation box after the interaction of a late endosomal membrane with **A.** a 70% OA PBAE particle **B.** a 30% OA PBAE particle **C.** an Onpattro – like LNP **D.** a 25 kDa bPEI polyplex **E.** a PPP polyplex. **F.** Lipids (excluding cholesterol) extracted from the plane of different membranes after interaction with the respective particles, mean ± sd, n = 3.

Based on umbrella sampling simulations, the octanol – water partition coefficients (log P) of the polymers were ranked according to their hydrophobicity: 70% OA PBAE (most hydrophobic) > PPP > 30% OA PBAE > bPEI (least hydrophobic) (Figure 4A+B). As discussed above, a direct correlation of polymer hydrophobicity with its EE performance can be seen as controversial. In this case, the most hydrophobic polymer caused the strongest membrane disturbance, but no clear relationship between the other polymers’ log P and their effect on endosomal membranes was found.

**Figure 4.**
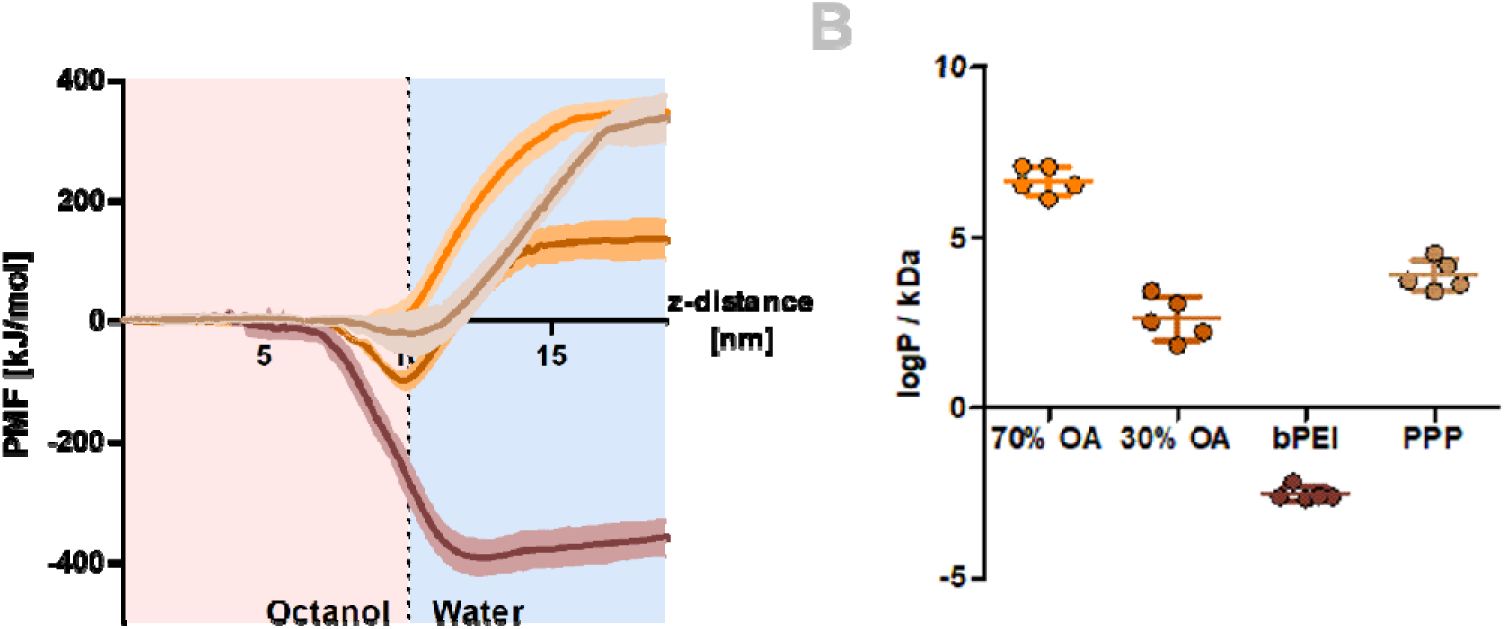
Determination of log P values in CG MD. **A.** Potential of mean force (PMF) curves of the transfer of a polymer molecule from an octanol phase to a water phase, mean ± sd (n = 5). **B.** log P values of polymers, normalized to molecular weight (per kDa) as calculated from PMF curves in A., mean ± sd (n = 5).

Secondly, electrostatic interactions were observed for all particles. As the nanoparticles contained an excess of cationic charges, the interaction with negatively charged membrane lipids was favorable. For the 70% OA PBAE, this correlated with the amount of lipids being extracted from the membranes (Fig. 3F). The least lipids were extracted from the least charged membrane type (lipid raft), and the most extraction took place from the strongly charged late endosome and lysosome membranes. The presence of glycolipids (late endosome vs. lysosome) however did not have a notable influence.

The preferred interaction of all polymers with negatively charged lipids was conclusive both in AA MD (Fig. 5A+B, Fig S6) and CG MD (Fig. 5C+D, Fig. S6+S7). In both cases, the anionic lipids (purple shades) clustered around the attached particle or free polymer molecules interacting with the membrane. Fig. 5 shows the interactions of 70% OA PBAE or bPEI with the late endosomal membrane. In AA resolution, the graphs are noisier due to the lower overall number of molecules in the simulations. Still, it is visible that both polymers (70% OA and bPEI) were transiently attached to the glycolipids, followed by favored contacts to other anionic lipids. This was in general true for all AA setups simulated (Fig. S6). The importance of anionic lipids for stable polymer–membrane interactions was particularly evident in simulations involving the lipid raft model. In this system, negative charges were confined to the glycosylated layer extending above the lipid headgroups. As a result, during the transient contacts with glycolipids no negative charges were available in the headgroup region of the membrane, and permanent polymer adsorption did not occur. The role of anionic lipids, especially the lysosome-specific lipid BMGP, has previously been discussed to be of high relevance for EE of cationic formulations/drugs^53, 54^ and was emphasized again by the here presented results. Additionally, the AA simulations confirmed the presence of interactions with neutral lipids and cholesterol for the PBAEs (Fig. 5B), which clearly distinguished them from the bPEI polymer.

**Figure 5.**
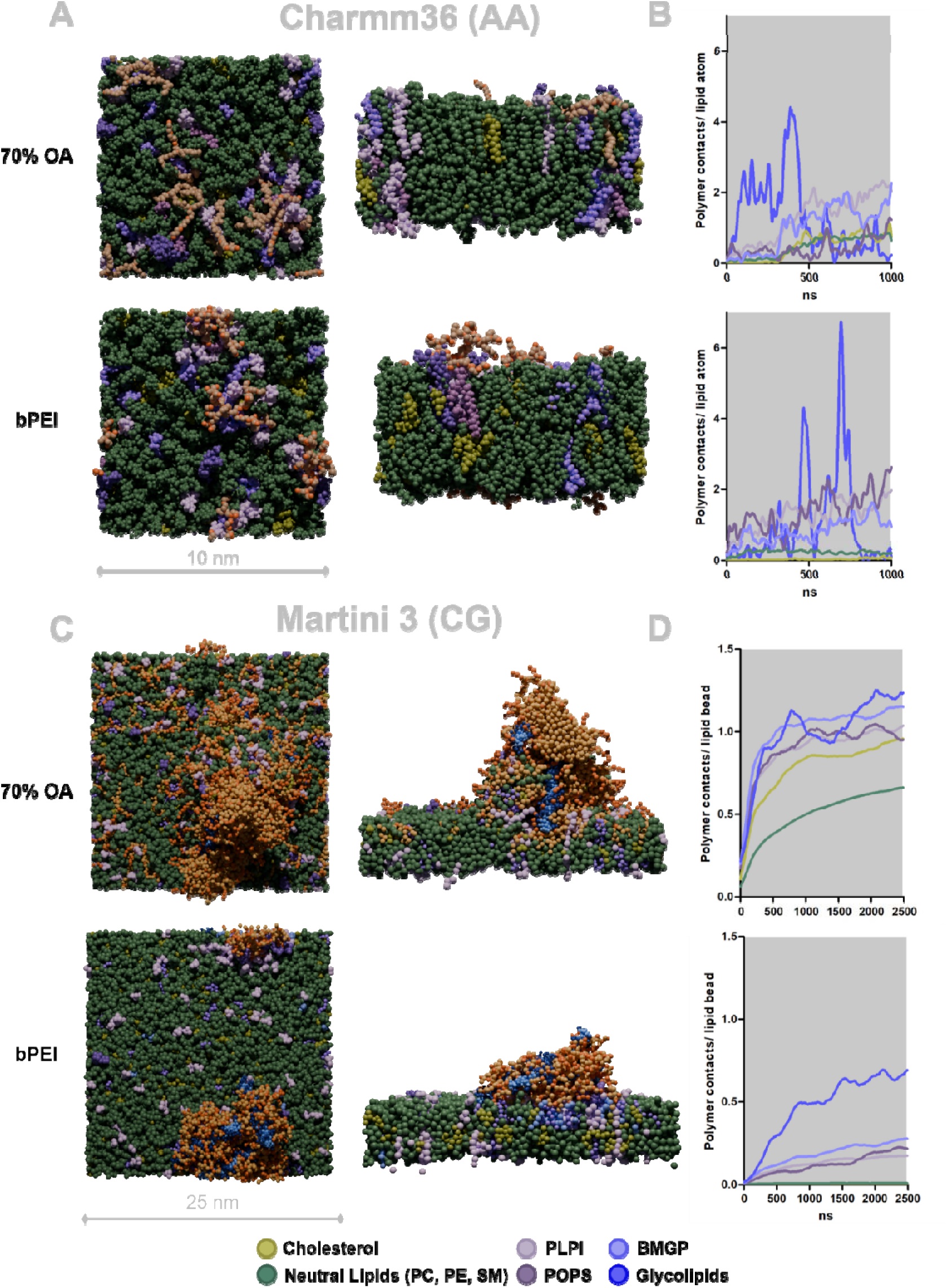
Contacts with late endosomal membrane lipids over time in AA and CG-MD simulations. **A.** Visualization of polymer – late endosome interactions (top- and side view) from AA simulations applying the charmm36 forcefield. **B.** Polymer contacts (upper: 70% OA PBAE, lower: bPEI) below 0.6 nm per membrane lipid atom (late endosome) over time, n = 2. **C.** Visualization of particle – late endosome interactions (top- and side view) from CG simulations in Martini 3. **D.** Polymer contacts (upper: 70% OA PBAE, lower: bPEI) below 0.6 nm per membrane lipid bead (late endosome) over time, n = 3.

Analysis of the CG simulations (Fig. 5C+D, Fig. S7+8) regarding polymer/MC3 – membrane contacts confirmed good agreement with the AA setups, although glycolipid contacts appeared less transient. In CG simulations, all particles remained associated with the membranes, including the lipid raft model, over the whole simulated timespan, likely due to the larger system size and hence an increased number of initial contact points.

### Interaction of nanoparticles with membrane vesicles

For further investigation of potential EE mechanisms, all particles were simulated in endosome-mimicking CG membrane vesicles (Fig. S9). The vesicles had varied inner diameters between 19 nm (lipid raft) and 24 nm (lysosome) to acknowledge the fact that the lysosome tends to be larger than the early endosome^55^. In the early endosome and the lipid raft, simulations were begun with 1 µs at pH 7.4 protonation settings and then continued with 7 µs at pH 6.5. In the late endosome and the lysosome, pH 7.4 was only simulated for the first 0.6 µs, followed by 1.2 µs at pH 6.5 and prolonged to 8 µs at pH 5.4 protonation. Similarly to the previous results, the affinity of the polymers and the ionizable lipid MC3 to negatively charged lipids was observed (Fig. S8) with increasing contacts over time. In general, the number of polymer contacts per lipid bead rapidly increased in the initial ns and then stabilized towards the end of the simulations.

The EE of lipid-based systems, such as the LNP incorporated in this study, is believed to rely on membrane fusion and the disruption of the membrane bilayer^56, 57^. In the vesicle simulation setups, the LNP fused with the membrane, which caused rapid exchange of lipids between membrane and LNP. Similarly as recently portrayed by others^36^, this caused disruption of the membranes and the formation of disordered phases in the vesicle (Fig. 6A+B). However, in only one of the eight LNP-vesicle interactions, this led to successful escape of siRNA molecules from the vesicle (Fig. 6A). Closer observation of this simulation revealed that the EE of two siRNA molecules in this simulation occurred directly during the initial fusion of LNP and vesicle. The energetic hurdle that must be overcame for the fusion of LNP and membrane to be initiated^52, 58^ seemed to be increased in the interaction with the lipid raft model. Here, no membrane fusion occurred (Fig. 6C) in one simulation, while in the repeated simulation fusion only occurred after ∼ 3 µs (Fig. S8). This can be explained by the absence of anionic lipids, as described above, or the presence of an increased amount of glycosylated lipids, forming a “buffer zone” above the membrane surface. The effect of pH in the endosomal compartments on the LNP-membrane interaction has been investigated in depth elsewhere^22^, with the result that acidic pH (< 6.5) enhances LNP disintegration and promotes lipid exchange between LNP and endosomal membranes. This was well reproduced by our results, as each step of pH reduction in the simulations led to an abrupt increase in the number of contacts between LNP lipids and membrane (Fig. S8).

**Figure 6.**
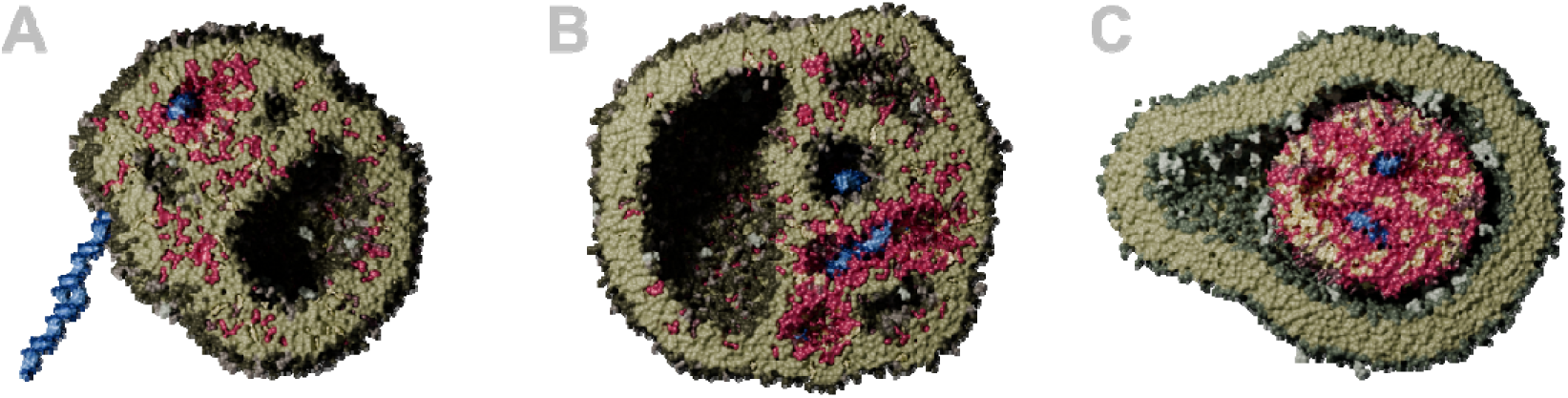
CG MD simulation outputs of an Onpattro® like LNP interacting with endosome mimicking vesicles for 8 µs. **A.** LNP and early endosomal vesicle. **B.** LNP and late endosomal vesicle. **C.** LNP and lipid raft vesicle. (green: membrane lipids, light gray: glycans, light green: cholesterol, blue: siRNA, pink: MC3/MC3H, purple: DSPC)

Interestingly, the 70% OA PBAE, even though to a smaller extent, caused membrane disturbances comparable to the LNP (Fig. 7A). This formulation preferably interacted with the anionic lipids but was at the same time capable of forming hydrophobic interactions with all membrane lipids (Fig. S8). Over time, the polyplex disassembled and polymer molecules distributed over the whole vesicle (Fig. 7A). In the lipid raft vesicle, the disintegration of the polyplex was less pronounced. Instead, incorporation of preferably cholesterol from the membrane into the hydrophobic PBAE core was observed (Fig. 7B). Transmission electron microscopy (TEM) imaging has previously been used to visualize the effect of EE efficient polyplexes on endosomes^8^. Those images showed disturbed membranes that could be interpreted similar to the results from the vesicle-70% OA PBAE polyplex simulations herein. However, no escape events were observed for the 70% OA PBAE polyplex. This may be attributed to the inherently low frequency of escape events, even in formulations considered effective for EE. Alternatively, the absence of detectable escape could suggest that PBAE-mediated EE involves a combination of membrane fusion and proton sponge-like mechanisms, which may have caused strong endosomal damage and Gal8 recruitment *in vitro*. In the vesicle simulations, chloride ions were inserted inside the vesicles to compensate for the increasing positive charge of the polymer under increasingly acidic conditions. However, this approach is unlikely to fully replicate the osmotic pressure dynamics that might develop physiologically.

**Figure 7.**
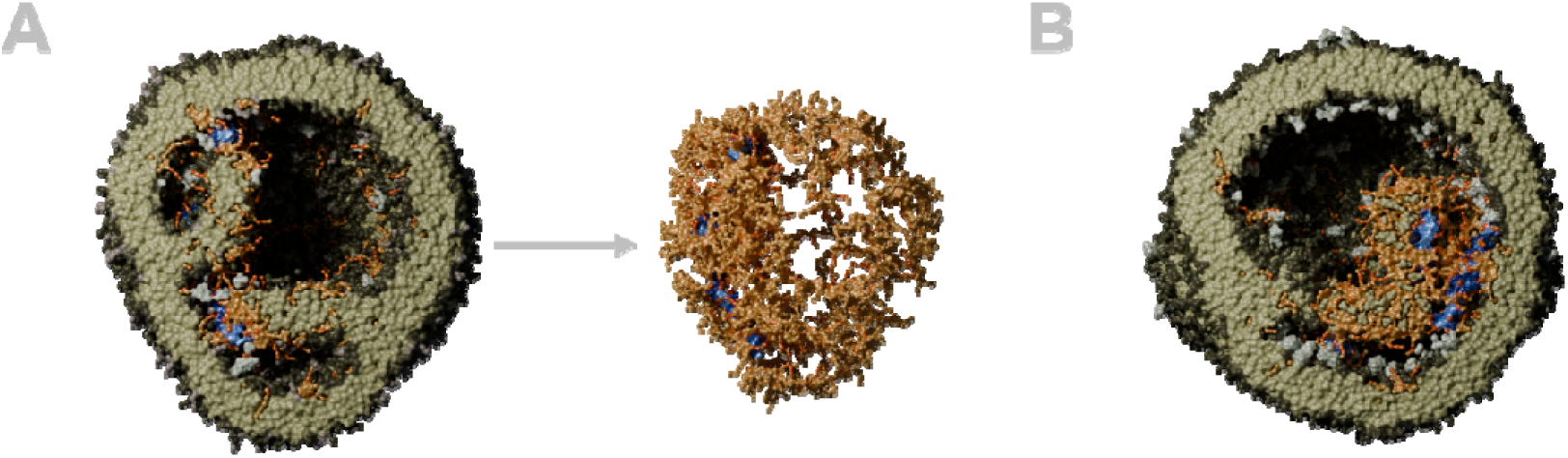
CG MD simulation outputs of 70% OA PBAE polyplex interacting with endosome mimicking vesicles for 8 µs. **A.** PBAE polyplex in the early endosomal vesicle (left), with visualization of disassembled polyplex only (right). **B.** 70% OA PBAE polyplex and lipid raft vesicle. (green: membrane lipids, light gray: glycans, blue: siRNA, beige/orange: polymer)

The more hydrophilic PBAE (30% OA) followed similar principles to the 70% OA PBAE, but due to the decreased hydrophobic interactions (Fig. S8), the effect of the particles on the vesicles was less pronounced (Fig. 8A+B). Instead of polymer molecules fusing into the membrane, the interaction was dominated by surface contacts between the cationic spermines and the anionic lipids. Only minor disturbances of the membrane bilayer occurred. However, the vesicles deformed to a flattened shape, which allowed more surface contact with the polyplex (Fig. 8B).

**Figure 8.**
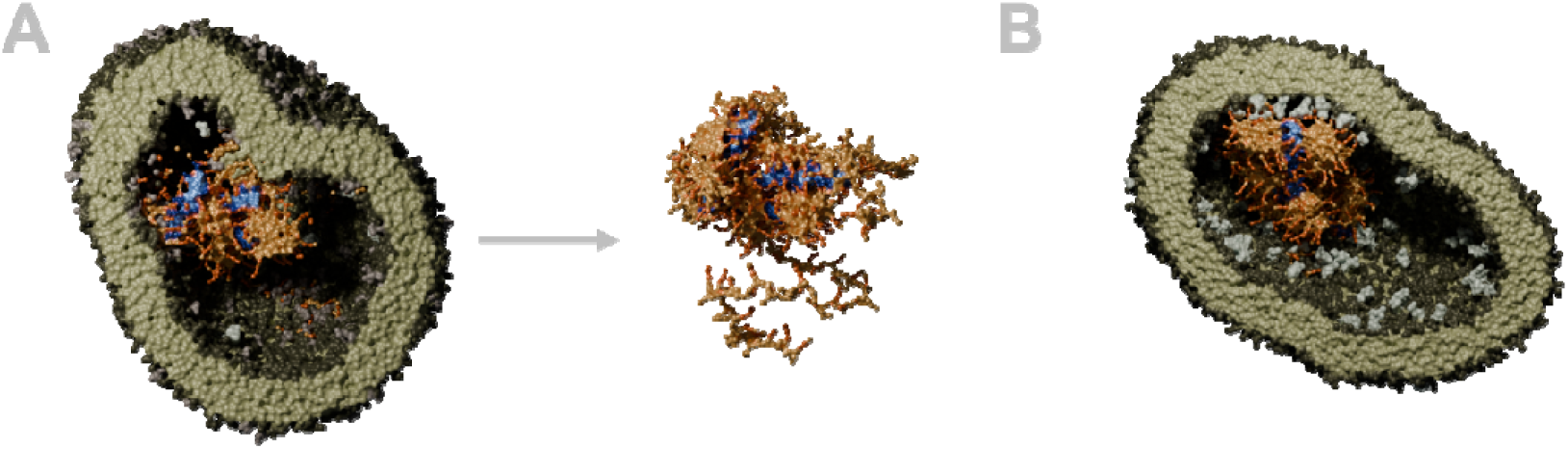
CG MD simulation outputs of 30% OA PBAE polyplex interacting with endosome mimicking vesicles for 8 µs. **A.** PBAE polyplex in the early endosomal vesicle (left), with visualization of disassembled polyplex only (right). **B.** 30% OA PBAE polyplex and lipid raft vesicle. (green: membrane lipids, light gray: glycans, blue: siRNA, beige/orange: polymer)

The effect of the bPEI polyplex on the membrane vesicles resembled the 30% OA PBAE, with the difference that the bPEI polyplex did not shed any polymer (Fig. 9A-C). As mentioned above, no hydrophobic interactions of the hydrophilic bPEI polymer with the membranes were observed (Fig. S8). The minimal membrane interaction observed for bPEI in this setup is consistent with its limited endosomal escape efficiency in the Gal8 recruitment assay. However, others have reported satisfying efficiency of PEI siRNA polyplexes before^33^. Some reported the formation of membrane pores due to the interaction with PEIs in simulation, however with the PEI being already placed in the membrane at the beginning of the simulation^59^. Neither our CG nor our AA models produced membrane pores in unstirred simulations. Only when the bPEI particle was not only pulled onto, but forcefully pulled through a membrane, a pore in the CG membrane formed (Fig. S10). In summary, this particle’s results support a hypothesis formed by others on the EE of PEI polyplexes in HeLa cells^15^: The highly charged PEI polyplex firmly associates with the membrane, and local osmotic or mechanical forces are necessary for RNA release through local membrane defects into the cytoplasm. As described above, the buildup of osmotic pressure in our unstirred equilibrium simulations is limited, which makes the observation of escape events in this setup unlikely.

**Figure 9.**
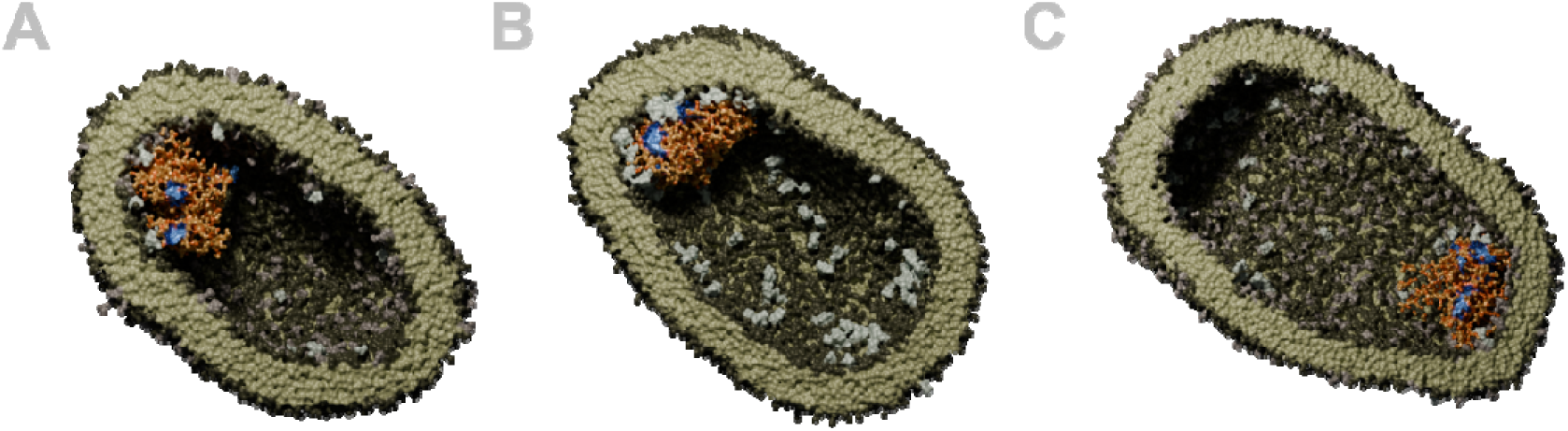
CG MD simulation outputs of the bPEI polyplex interacting with endosome mimicking vesicles for 8 µs. **A.** bPEI polyplex in the early endosomal vesicle **B.** bPEI polyplex in the lipid raft vesicle. **C.** bPEI polyplex in the late endosome vesicle. (green: membrane lipids, light gray: glycans, blue: siRNA, beige/orange: polymer)

Finally, the PPP polyplex performed comparably to the bPEI and the 30% OA particles (Fig. 10A+B). As described for the planar membrane interaction, the PCL segment accumulated in the lipid tail region of the membranes, but only if the particle did not attach to the membrane surface with the bPEI residues first. In this case, the particle was hindered from hydrophobic interactions as the PEI stuck to the anionic membrane surface (Fig. 10A). To investigate whether a larger PPP particle with therefore more hydrophobic units would cause membrane disturbance akin to the 70% OA polyplex, a PPP particle with a total polymer mass equal to the 70% OA particle was created. This polyplex caused some disturbance in the lysosomal membrane, but like the smaller particles, it did not disassemble (Fig. 10C).

**Figure 10.**
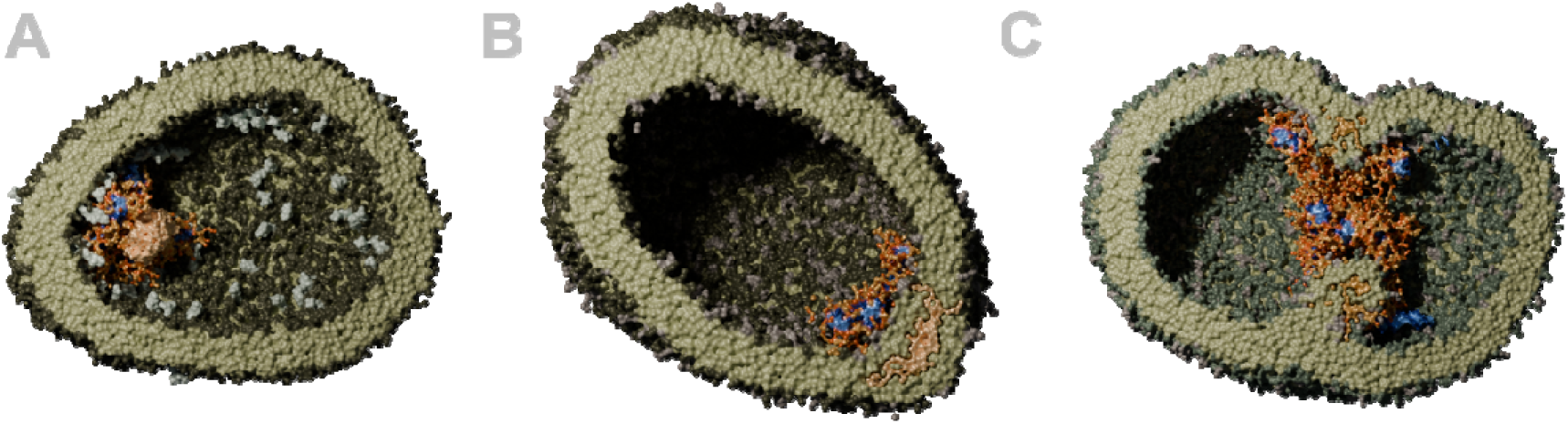
CG MD simulation outputs of the PPP polyplex interacting with endosome mimicking vesicles. **A.** Small PPP polyplex in the lipid raft vesicle **B.** Small PPP polyplex in the lysosomal vesicle. **C.** Larger PPP polyplex (containing 15 siRNA molecules) in the lysosomal vesicle. (green: membrane lipids, light gray: glycans, blue: siRNA, beige/orange: polymer (darker orange bPEI, PCL shown in brighter coloring)

The outcome of particle–vesicle interactions is summarized in Fig. 11A, reporting the total number of contacts between polymer (or MC3 plus DPSC for the LNP) and membrane lipids in the final simulation frame. Consistent with the observations from planar membrane simulations, both the LNP and the 70% OA PBAE polyplex exhibited substantially higher membrane contact numbers compared to the other formulations. Specifically, the total number of contacts between 70% OA PBAE and membrane lipids was between 18.6-fold (lysosomal membrane) and 52.8-fold (early endosomal membrane) greater than that observed for bPEI. Hence, the difference can not solely be attributed to the overall higher amount of polymer in the PBAE polyplex, but also polymer specific properties. Furthermore, as noted previously, the presence of anionic lipids appeared to promote interaction, with late endosomal and lysosomal vesicles showing higher abundance of polymer or LNP contacts than early endosomal or lipid raft membranes.

**Figure 11.**
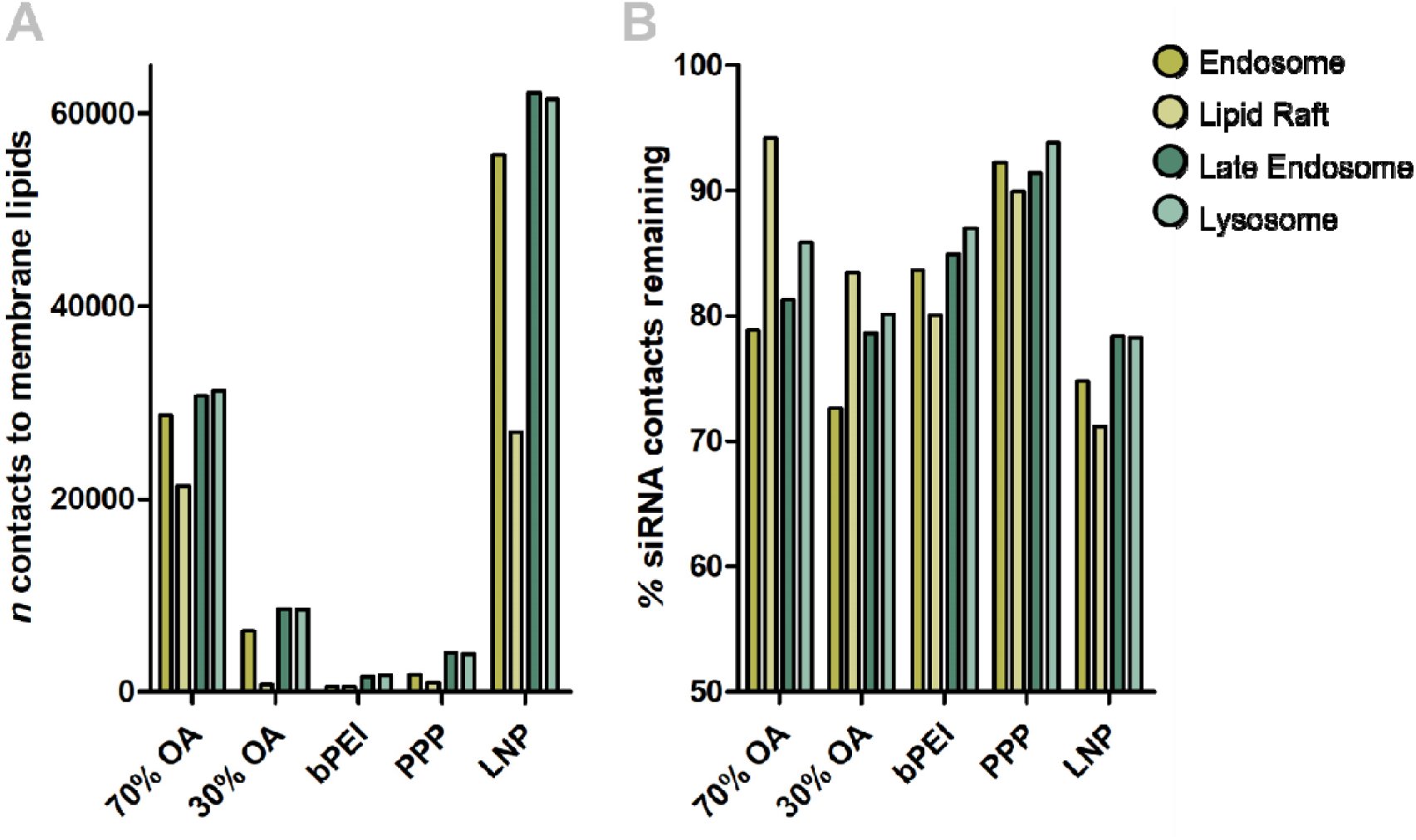
Quantification of CG MD output after 8 µs simulated interaction between endosomal membranes and model particles. **A.** Total number of contacts of the polymer (or DLin-MC3-DMG and DSPC for the LNP) with membrane components in the last frame of simulations (8 µs); shown as mean from n = 2 simulations per setup. **B.** Percentage of contacts of the polymer (or DLin-MC3-DMG and DSPC for the LNP) with siRNA that remained after 8 µs simulated interaction with model vesicles in comparison to t = 0 ns; shown as mean from n = 2 simulations per setup.

Polyplex stability is known to depend on multiple interrelated factors, including polymer architecture, molecular weight, and hydrophobicity^35, 60^. In an effective polyplex formulation, these factors must be well balanced to ensure sufficient stability in serum and efficient cargo release following cellular uptake^61^. In our CG MD simulations, a comparison of polymer–siRNA contacts before and after endosomal membrane interaction revealed a reduction in siRNA encapsulation across all systems tested (Fig. 11B), suggesting partial unpacking upon membrane contact. While no consistent trend emerged across membrane compositions, the results indicate that the 30% OA PBAE polyplex is the least stable, whereas the PPP-based polyplex shows the highest stability. Interestingly, the polyplex that exhibited the strongest membrane interaction (70% OA PBAE) did not show the most efficient siRNA unpacking. Nevertheless, comparisons between structurally related polymers (70% OA vs. 30% OA PBAE; PPP vs. bPEI) revealed that increased hydrophobicity was associated with reduced unpacking.

Noticeably, the MD-based unpacking results reproduced the outcome from an experimental stability assay (Fig. S11), in which competitive displacement by heparin and Triton X revealed an identical stability ranking among the four polyplexes. This correlation validates the predictive power of the simulation approach and highlights its ability to mechanistically dissect complex structure–function relationships.

## Conclusion

The herein presented MD simulations visualized and quantified the interaction of different siRNA nanoparticles with endo-lysosomal membranes. All formulations showed increased affinity to anionic membrane lipids, highlighting that these play a significant role for EE. However, only two formulations (the 70% OA PBAE polyplex and the Onpattro®-like LNP) achieved eGFP knockdown *in vitro*. These were the only particles that formed larger numbers of hydrophobic interactions in MD, extracting lipids from planar membranes and disturbing membrane vesicles. Hence, a pronounced role of hydrophobic interactions for effective EE through membrane fusion was demonstrated. Particles that do not contain hydrophobic residues (such as bPEI polyplexes) were not capable of interacting with the lipid tails of endosomal membranes.

Combining the findings of this work with the existing knowledge on EE mechanisms, we understand EE mechanisms of polyplexes as a spectrum ranging from proton sponge-like EE to membrane fusion-dominated mechanisms. For the EE of polyplexes based on hydrophilic polymers such as PEI, our simulations propose an electrostatically driven attachment of particles onto endosomal membranes, which could be followed by comparably rare escape events through the buildup of osmotic or mechanical stress. Larger hydrophobic modifications of a polymer are necessary for hydrophobic interactions to arise, which enhances membrane disruption and polymer shedding. By exploiting both proton-sponge like and membrane fusion-based mechanisms, the amphiphilic polyplex formulations (represented herein by 70% OA PBAE polyplexes) possess increased EE efficiency, but also cause large membrane defects, visible *in vitro* through the observation of Gal8 recruitment. This type of polyplex is efficient at siRNA delivery, but potentially cytotoxic through excessive endosomal disruption.

In our simulations, the Onpattro®-like LNPs caused the highest extent of membrane disruption through fusion of LNP and membrane and the mixing of lipids. *In vitro*, the formulation produced only small holes in the endosome, which were not detectable by galectins. Considering the cytotoxicity caused by large endosomal defects, the EE mechanism of the simulated LNP appears desirable. Engineering polyplexes towards more membrane fusion-based EE, for example by modifying hydrophobic residues, could therefore largely advance the development of polymers for siRNA delivery.

Even though escape events of siRNA from polyplex formulations were not observed in simulation, our results show that MD simulations of EE mechanisms can conclusively be correlated to *in vitro* results. In the future, expanding the library of simulated nanoparticle-membrane systems will help to solidify the identification of desirable EE mechanisms in MD simulations. Subsequently, MD could be used to predict EE behavior of a formulation and screen for beneficial properties.

## Materials and Methods

### Materials

Spermine and oleylamine for the synthesis of the poly(beta)aminoesters (PBAE) were obtained from Fisher Scientific (Acros, USA), whereas the 1,4-butanediol diacrylate for the PBAE backbone originated from Tokyo Chemical Industry Co. (Tokyo, Japan). The 25 kDa branched polyethylenimine (bPEI) and the 5 kDa bPEI for the synthesis of the PEI-PCL-PEI polymer (PPP) were a kind gift from BASF (Ludwigshafen, Germany). Polycaprolactone diacrylate (PCL), as well as HEPES (4-(2-hydroxylethyl)-1-piperazineethanesulfonic acid), Dulbecco’s Phosphate Buffered Saline (PBS), D-Glucose, Triton-X, porcine Heparin sodium salt, Cholesterol and Copper sulfate pentahydrate, Minimum Essential Medium Eagle (MEM), High/Low glucose Dulbecco’s Modified Eagle Medium (DMEM), fetal bovine serum (FBS), MEM Non-Essential Amino Acids (NEAA) solution (100X) and Cell Counting Kit-8 were obtained from Sigma-Aldrich (Taufkirchen, Germany). D-Lin-MC3-DMA was from MedChemExpress (Sollentuna, Sweden), DSPC and DMG-PEG 2000 from Avanti Research (Birmingham, United Kingdom). The siRNA used in this project was amine-modified siRNA for the knockdown of eGFP (siGFP), with the sequence 5’-pACCCUGAAGUUCAUCUGCACCACcg, 3’-ACUGGGACUUCAAGUAGACGGGUGGC, or negative control siRNA (siNC) with the sequence 5’-pCGUUAAUCGCGUAUAAUACGCGUAT, 3’-CAGCAAUUAGCGCAUAUUAUGCGCAUA, both purchased from Merck (Darmstadt, Germany). Dyes for confocal microscopy (DAPI (4’,6-Diamidin-2-phenylindol, Dihydrochloride), Alexa-Fluor 647) were obtained from Life Technologies GmbH (Frankfurt, Germany). Penicillin-Streptomycin (P/S, 10.000 U/ml), Blasticidin S HCl (10 mg/ml), Lipofectamine 2000, 4% Paraformaldehyde in PBS and DAPI (4′,6-diamidino-2-phenylindole) were bought from Thermo Fisher (Waltham, MA, USA). CytoTox 96® Non-Radioactive Cytotoxicity Assay kit was purchased from Promega Corporation (Madison, WI, USA)

### Experimental Methods

#### Polymer Synthesis

The poly(beta)aminoesters (PBAE) were synthesized via Michael Addition as previously described^35^. The synthesis was based on tri-boc-spermine as hydrophilic side chain, oleylamine (OA) as hydrophobic side chain, and 1,4-butanendiol diacrylate as backbone. After polymerization, the tri-boc-spermine was deprotected with trifluoroacetic acid. The ratio of hydrophilic spermine to hydrophobic OA was controlled by the input ratio of the reagents and validated by ^1^H nuclear magnetic resonance (NMR). For this work, a polymer containing 33% OA (generally referenced as “30% OA” for clarity in the comparison to simulation) and 68% OA (“70% OA”) were used.

The synthesis of the 5 kDa-5 kDa-5 kDa PEI-PCL-PEI polymer (PPP) was previously described by Jin et al.^34^ In brief, 5 kDa branched polyethylenimine (bPEI) was stirred with polycaprolactone-diacrylate (PCL) for 48 h at 40 °C at a molar ratio of 2 (PEI) : 1 (PCL). Then the polymer was purified from monomers by dialysis against water with a molecular weight cut-off of 10 kDa and subsequently lyophilized. The ratio of PEI:PCL in the product was evaluated to be 2:1 by a TNBS assay, which was calibrated with unmodified 5 kDa PEI^62^.

#### Particle Formulation

##### PBAE and bPEI Particles

PBAE and 25 kDa bPEI polyplexes were formulated at an N/P ratio of ten (i.e., ten protonable units of the polymer per phosphate of the siRNA). The polymers and the siRNA were separately diluted in 10 mM HEPES buffer at pH 5.4 to equal volumes. For the particles used in the hemolysis assay, HEPES-buffered Glucose (5%) (HBG) at three different pH levels (7.4, 6.5 and 5.4) was used instead to ensure isotonicity. Both components were mixed by batch-mixing and shortly vortexed afterwards. Finally, the particles were incubated for 90 minutes at room temperature before further use.

##### PPP Particles

The PPP particles were prepared as previously described^34^ by preparing empty particles first and subsequently adding the siRNA. The polymer was dissolved in acetone, and then slowly dripped into formulation buffer (as above) while stirring. The acetone was left to evaporate from the mixture over three hours, and the formation of empty particles was confirmed by dynamic light scattering (DLS). Then, siRNA loaded polyplexes were prepared by diluting the empty particles and mixing as described above for the PBAE and bPEI particles.

##### MC3 – LNP

For the preparation of LNPs, the lipids were diluted in ethanol, with ratios of 50% Dlin-MC3-DMA, 38.5% cholesterol, 10% DSPC and 1.5% PEG-2000-DMG. The siRNA was diluted in 25 mM sodium acetate buffer at pH 4. Lipid blend and siRNA were combined by microfluidic mixing in a T-mixer (micro IDEX H&S P-888) at flow rate ratios of 0.75 ml/min (lipid) and 2,25 ml/min (siRNA). Afterwards, the LNPs were dialyzed overnight against 150 mM PBS or HBG when used for the hemolysis assay. Before further use, the LNPs were filtered through a 0.22 µm syringe filter.

Z-average, PDI and ζ-potential of all particles were determined on a Malvern Zetasizer Ultra (Malvern Instruments, Malvern, UK). All particles were produced in biological triplicates for characterization.

#### Hemolysis Assay

For the hemolysis assay^63^, blood from a healthy anonymous donor was centrifuged for 10 minutes at 1500x g. The sedimented erythrocytes were resuspended in PBS and repeatedly washed with PBS until the supernatant was clear after centrifugation. The erythrocytes were resuspended in PBS again and then diluted to 5 * 10^8/ml in HBG with either pH 5.4, 6.5 or 7.4. Subsequently, 1:1 mixtures of erythrocytes and the respective particles (50 pmol siRNA/ 100 µl) or 15 mg/ml polymer dilutions were then incubated in 96-well plates for 30 minutes at 37 °C. 1% Triton-X was used as positive control (i.e., 100% hemolysis) and buffers as negative controls. After incubation, the plates were centrifuged for 5 minutes at 1500x g and the supernatants were transferred to fresh transparent well plates. The absorbance of the supernatants was measured at 541 nm on a TECAN Spark Plate Reader (Tecan Trading AG, Switzerland). The assay was performed in a biological triplicate. Erythrocyte aggregation was documented on the resuspended erythrocyte pellets on a Evos M5000 microscope (Thermo Fisher Scientific, Schwerte, Germany).

#### Particle Stability Assay

The stability of polyplex formulations was assessed by a competition assay at pH 5.4, as previously described elsewhere^35^. Briefly, a dilution series of stress solutions containing heparin and Triton X was prepared. In this regard, 100% stress referred to a concentration of 200 USP units heparin/ml and 1% (m/m) Triton X. First, 10 µL of nanoparticles formulated at pH 5.4 were incubated with 20 µL of the respective stress solution for 1 h at 37 °C. Then, 5 µL of diluted SYBR Gold solution was added and after 5 minutes, fluorescence was measured on a TECAN Spark Plate Reader (Tecan Trading AG, Switzerland). Excitation was set to 492 nm; emission was set to 537 nm. For data analysis, all data points (n = 3 technical replicates) were normalized towards the 100% stress sample of each formulation, which was assumed to represent maximal unpacking (i.e., 0% encapsulation). To calculate the EC50 value of each formulation, a sigmoidal curve fit with automated outlier detection was performed in GraphPad Prism5 2007 software.

#### Cell Culture

Hela WT cells (passages 10-15) were cultured in MEM containing 10% FBS, Hela/eGFP cells (ATCC, USA, passages 5-10) were cultured in DMEM-high glucose containing 10% FBS, 0.1 mM MEM NEAA, 1% P/S and 10 μg/mL Blasticidin. Hela-Gal8-mRuby3 cells (passages 5-10) were kindly provided by Professor Ernst Wagner (Ludwig-Maximilians-Universität Munich, Germany) and cultured in DMEM-low glucose with 10% FBS and 1% P/S. All cells were cultured in a humidified atmosphere containing 5% CO_2_ at 37 °C.

#### Galectin-8 Assay

The recruitment of Galectin-8 (Gal8) to damaged endosomes as an indicator of successful endosomal escape was tested on HeLa cells expressing mRuby-3-Gal8 fusion protein. Cells were seeded at a density of 10,000 cells per well in an 8-well ibiTreat chamber slide (Ibidi, Gräfelfing, Germany). The cells were transfected with nanoparticles containing 20 pmol siRNA per well for 4 hours or 24 hours. Of the total siRNA, 20% were labeled with Alexa Fluor 647. The culture medium was changed after 4 hours, and cells were imaged at the SP8 inverted confocal laser scanning microscope (CLSM; Leica Camera, Wetzlar, Germany) with a 63X oil objective. Cell nuclei were stained with DAPI. siRNA uptake and Gal8 puncta of ≥ 25 cells per sample were quantified from the images by automated counting using the Fuji plug-in of Image J.

#### eGFP Knockdown

HeLa/eGFP cells were seeded at a density of 6,000 cells per well in 96-well plates. The following day, cells were transfected with nanoparticles containing either 20 pmol siRNA targeting eGFP mRNA (siGFP) or 20 pmol scrambled siRNA of the same length (siNC) for 48 hours. Lipofectamine 2000 was used as a positive control, while free siRNA served as a negative control. After incubation, the cells were collected to perform the FACS analysis (Attune NxT Flow Cytometer, ThermoFisher Scientific). The eGFP knockdown efficiencies (biological duplicate with n = 3 technical replicates) were calculated by dividing the Median Fluorescence Intensity (MFI) of siRNA-treated group by that of the respective siNC-treated group.

#### Cellular uptake

HeLa/eGFP cells were seeded at a density of 6,000 cells per well in 96-well plates. The following day, cells were transfected with nanoparticles containing 20 pmol siGFP, of which 20% were labelled with Alexa Fluor 647. Lipofectamine 2000 was used as a positive control, while free siRNA served as a negative control. After 24 hours, the cells were collected to perform the FACS analysis (Attune NxT Flow Cytometer, ThermoFisher Scientific).

#### LDH- and CCK-8 Assay

To evaluate cytotoxicity, both an LDH- and a CCK-8 assay were conducted (n = 3 technical replicates). The particles for this experiment were prepared as described above and then diluted to test four concentrations (i.e., 40, 30, 20 and 10 pmol siRNA/50 µL). HeLa cells were seeded at 6,000 cells in 96 well plates. When reached 80% confluence, the cells were incubated with the four nanoparticle dilutions for 48 hours. For the LDH-assay, 50 µL cell culture supernatant was diluted 1:1 with CytoTox 96 reagent and incubated in the dark for 30 minutes at room temperature. After addition of 50 µL stop solution, the samples were quantified at 490 nm absorbance, and the results were normalized to the positive control. The cells pre-treated with 20 µL of lysis solution for 45 minutes at 37°C served as positive control for maximum LDH release. For the CCK-8 assay, 10 µL CCK-8 solution was added directly into each well containing treated cells and incubated for another 3 hours. The cell viability was quantified by absorbance relative to an untreated sample at 450 nm.

### CG-MD Simulation

All simulations were run in Gromacs 2021.4-plumed^64^. For the CG simulations, the Martini 3 forcefield^30^ was applied. The CG model particles were generated with our previously established siRNA model^31^ and contain three siRNA molecules each. The number of polymer molecules was chosen to reach ∼ N/P 10 for all polyplexes, whereas the MC3 – LNP had a final N/P ratio of 6.5. After minimization and NPT equilibration, all simulations were run at a timestep of 15 fs with Particle mesh Ewald (PME) electrostatic handling^65^ with a cutoff of 1.1 nm. Temperature was controlled by v-rescale temperature coupling at 298 K (particle assembly) or 310 K (membrane interactions), whereas pressure was handled by the Parrinello-Rahman barostat^66^ at 1 bar.

#### CG Particle Models

##### PBAE and bPEI Particles

The PBAE model particles were generated in accordance with our previously established approach for this group of polymers^31^ with either a 30% Oleylamine (OA) or a 70% OA polymer model. The bPEI particles were generated the same way, based on a 25 kDa PEI model with a branching degree of 59%^67^ (bPEI). PBAE or bPEI particles initially self-assembled in a 5 µs unbiased run in a cubic box with 26 nm side length. Boxes were solvated with 10 mM HEPES and neutralized with chloride ions. During the first run, the protonation corresponded to pH 5.4. This run was followed by 0.5 µs run time with reduced polymer protonation, allowing the particles to adjust to a theoretical pH 7.4. To generate input for the interaction with planar membranes at pH 6.5 or 5.4, the protonation was subsequently adjusted for another 0.5 µs. All particles were extracted from their initial assembly box including the ions of their hydration shell to ensure the particle stability in new simulation boxes with PME electrostatics.

##### PPP Particles

The PPP polymer model was created from two identical 5 kDa bPEI models^67^ that were connected by a 5 kDa PCL chain. The parametrization of this PCL chain was obtained according to the common Martini parametrization approach (*cgmartini.nl/index.php*)^68–71^. To ensure a particle constitution comparable to wet-lab experiments, the PPP polymer was left to self-assemble for 3 µs under the simulation conditions described above. The siRNA molecules were then added to the empty particle in a subsequent 2.5 µs simulation for encapsulation. pH adjustments were done in the same way as for the PBAE and bPEI particles.

##### MC3 – LNP

The MC3 – LNP with an Onpattro® – like composition was assembled following the recently published approach by Kjølbe et al.^36^ The siRNA is placed in water channels within a hexagonal lipid core, covered by an outer lipid layer. This resulted in a particle consisting of 49% MC3, 39% Cholesterol and 12% DSPC and a total N/P ratio of ∼ 6.5. PEG-lipids were not incorporated, as they are expected to be already shed from the LNP when it reaches the endosomal compartment of cells^38^. Protonation of MC3 was calculated based on the apparent pKa of 6.55 for MC3^72^, however the lipids in direct contact with the siRNA were kept protonated at all pH levels studied to avoid excessive shedding of siRNA from the LNP at pH 7.4.

All particle models and the chemical structures of the polymers are depicted in Fig. 1.

#### Membrane Models

To study the interaction of nanoparticles with different membrane compositions present in the endo-lysosomal pathway, four different membrane types were established, namely an early endosomal membrane, a lipid raft, a late endosomal membrane and a lysosomal membrane. The respective compositions were adapted from literature^50, 73–77^ and are listed in table S1. The initial topologies were generated with the INSANE^78^ tool. Lipid topologies were used from literature^30, 79, 80^ or created based on the established building blocks if not yet available in Martini 3 (e.g., bis(monoacylglycero)phosphate (BMGP), glycolipids (DPG1 and DPG3^79, 81^)). All planar models had an initial size of 25×25 nm and were simulated with semi-isotropic pressure scaling. The membranes were fixed in their position by restraints on the z-coordinate applied to the POPC molecules (in case of the lipid raft, only 50% of POPC molecules) with a force constant of 1000 kJ/mol*nm^2^. Lateral diffusion of lipid components (POPC, DPSM or cholesterol) inside the planar models was analyzed with the mean squared displacement tool^82, 83^ from the MDAnalysis package^84, 85^. For the vesicular models, membrane discs with a radius of 25 nm (early endosome), 26 nm (lipid raft) or 28 nm (late endosome, lysosome) were generated. To initiate the formation of vesicles, the membranes’ center of mass (COM) was pulled out of the plane in z-direction by a moving restraint applied with the PLUMED plugin. Afterwards, closed vesicles formed in an unbiased simulation within less than 1 µs.

#### Umbrella Sampling

The lipophilicity of the simulated CG polymer models was characterized by the calculation of a logP/ kDa according to eq. 1:

(1)

The free energy ΔG_w/o_ of transferring one polymer molecule from an octanol phase to a water phase was determined in umbrella sampling simulations^86, 87^. A polymer molecule and neutralizing chloride ions were placed in the center of an octanol phase of 9x9x20 nm. Then the molecule was pulled into the adjacent water phase with equal dimensions by a harmonic potential applied along the z-axis. A spacing of 0.25 nm was used for the umbrella windows, resulting in 65 windows, which were simulated for 20 ns each. The potential of mean force (PMF) was generated as a function of distance from initial position by weighted histogram analysis with the gmx *wham* function^88–90^.

#### Planar Membrane Interactions

For the investigation of particle-membrane interactions in a simple, planar setup, the 25×25 nm membranes were simulated with each model particle (70% OA PBAE, 30% OA PBAE, bPEI, PPP and LNP) in triplicate runs. The box had a z-dimension of 35 nm, and the membranes were centered at z = 10 nm. All boxes were solvated with 150 mM NaCl and simulated at 310 K. The interactions with the early endosome and the lipid raft were simulated at pH 6.5, whereas late endosome and lysosomal interaction was simulated with protonation settings of the particles corresponding to pH 5.4. To ensure interaction from the glycosylated membrane leaflet, the particles were pulled in contact with the membrane by a moving restraint with the PLUMED plugin, lasting for 5 ns. Then a 2495 ns unbiased interaction was simulated with the above-mentioned settings. For analysis, the mass density distribution along the z-axis was calculated vie the gmx *density* function, defining the center of the membrane as z = 0. The amount of membrane components extracted from the membrane plane was calculated from the density distribution, where the upper limit of the membrane area was defined as the point, where the first derivative of the density distribution was > −100.

#### Vesicle Interactions

Additionally, the interaction of particles with the inside of endosome-mimicking vesicles was simulated to compare a more realistic setup. Particles were placed inside the preformed vesicles and solvated with 150 mM NaCl at 310 K in cubic boxes with 40 or 42 nm side length. All setups started with protonation corresponding to pH 7.4. For the early endosome and the lipid raft, the pH was reduced to 6.5 after 1 µs, which was then followed by 7 µs simulation at the mildly acidic pH. In the late endosome and the lysosome vesicle, pH was reduced to 6.5 after 0.5 µs, and then further reduced to pH 5.4 after another 1.2 µs, at which additional 6.3 µs were simulated. The additional chloride ions needed to neutralize the polymer charge after each pH change were placed inside the vesicle to mimic the osmotic pressure increase inside the endosome. Overall, every vesicle-particle setup was simulated for 8 µs and every setup was simulated twice. The vesicle interactions were analyzed for the number of polymer–membrane contacts over time using the gmx *mindist* tool and for changes in the siRNA environment via the Radial Distribution Function (RDF).

### AA Models and Simulations

The AA simulation input was generated with CHARMM-GUI^91, 92^ using the CHARMM36 forcefield^93^: Planar membrane models with a size of 10×10 nm and lipid compositions identical to the CG membranes were built by the Membrane builder^94–96^. Polymer models with a reduced molecular weight (trimers for PBAEs, i.e. ∼ 1.3 kDa; bPEI and PPP ∼ 1.8 kDa) were obtained through the Ligand Reader & Modeler^97^. All AA simulation boxes were neutralized and solvated with 150 mM sodium chloride. The simulations were run as NPT ensembles with the standard settings supplied by CHARMM, i.e. PME electrostatic handling, semi-isotropic pressure scaling with the c-rescale barostat, v-rescale thermostat at 310.15 K and a timestep of 2 fs. The membranes were first simulated for 500 ns without interacting polymers. For the simulation of polymer-membrane interactions, ∼ 9 kDa total mass of a polymer were used. The PBAEs were simulated separately for 500 ns to form micelles similar as the CG model and then further used in this form. The bPEI and the PPP models were used as individual molecules, as they did not aggregate within 500 ns (for PPP unlike the CG model – arguably due to the reduced MW and therefore length of the PCL segment). All polymers were placed in proximity to the glycosylated membrane leaflets in a 10×10×18 nm box. To facilitate initial contact with the membrane, a short pulling sequence mediated by a PLUMED moving restraint was applied to the polymer molecules. The polymer-membrane interactions were simulated in duplicates for 1 µs and analyzed with the gmx *mindist* tool.

### Data Analysis and Visualization

Graphs were created in GraphPad Prism5 2007 software, which was also used for statistical analysis, where applicable. Simulations were visualized in Blender 4.5.2 LTS.

## Acknowledgments

The authors gratefully acknowledge the European Research Council (ERC-2022-COG-101088587) for funding this project and the Gauss Centre for Supercomputing e.V. (www.gauss centre.eu) for providing computing time on the GCS Supercomputer SUPERMUC-NG at Leibniz Supercomputing Centre (www.lrz.de) in the framework of the Project “AVOCADO2”.

## References

1. Jadhav, V.; Vaishnaw, A.; Fitzgerald, K.; Maier, M. A., RNA interference in the era of nucleic acid therapeutics. Nature Biotechnology 2024, 42 (3), 394–405.

2. Whitehead, K. A.; Langer, R.; Anderson, D. G., Knocking down barriers: advances in siRNA delivery. Nat Rev Drug Discov 2009, 8 (2), 129–38.

3. Jeon, T.; Luther, D. C.; Goswami, R.; Bell, C.; Nagaraj, H.; Cicek, Y. A.; Huang, R.; Mas-Rosario, J. A.; Elia, J. L.; Im, J.; Lee, Y.-W.; Liu, Y.; Scaletti, F.; Farkas, M. E.; Mager, J.; Rotello, V. M., Engineered Polymer–siRNA Polyplexes Provide Effective Treatment of Lung Inflammation. ACS Nano 2023, 17 (5), 4315–4326.

4. El Moukhtari, S. H.; Garbayo, E.; Amundarain, A.; Pascual-Gil, S.; Carrasco-León, A.; Prosper, F.; Agirre, X.; Blanco-Prieto, M. J., Lipid nanoparticles for siRNA delivery in cancer treatment. Journal of Controlled Release 2023, 361, 130–146.

5. Schiffelers, R. M., Cancer siRNA therapy by tumor selective delivery with ligand-targeted sterically stabilized nanoparticle. Nucleic Acids Research 2004, 32 (19), e149–e149.

6. Urban-Klein, B.; Werth, S.; Abuharbeid, S.; Czubayko, F.; Aigner, A., RNAi-mediated gene-targeting through systemic application of polyethylenimine (PEI)-complexed siRNA in vivo. Gene Ther 2005, 12 (5), 461–6.

7. Vermeulen, L. M. P.; Brans, T.; Samal, S. K.; Dubruel, P.; Demeester, J.; De Smedt, S. C.; Remaut, K.; Braeckmans, K., Endosomal Size and Membrane Leakiness Influence Proton Sponge-Based Rupture of Endosomal Vesicles. ACS Nano 2018, 12 (3), 2332–2345.

8. Kilchrist, K. V.; Dimobi, S. C.; Jackson, M. A.; Evans, B. C.; Werfel, T. A.; Dailing, E. A.; Bedingfield, S. K.; Kelly, I. B.; Duvall, C. L., Gal8 Visualization of Endosome Disruption Predicts Carrier-Mediated Biologic Drug Intracellular Bioavailability. ACS Nano 2019, 13 (2), 1136–1152.

9. Jin, Y.; Wang, X.; Kromer, A. P. E.; Müller, J. T.; Zimmermann, C.; Xu, Z.; Hartschuh, A.; Adams, F.; Merkel, O. M., Role of Hydrophobic Modification in Spermine-Based Poly(β-amino ester)s for siRNA Delivery and Their Spray-Dried Powders for Inhalation and Improved Storage. Biomacromolecules 2024, 25 (7), 4177–4191.

10. Werfel, T. A.; Jackson, M. A.; Kavanaugh, T. E.; Kirkbride, K. C.; Miteva, M.; Giorgio, T. D.; Duvall, C., Combinatorial optimization of PEG architecture and hydrophobic content improves ternary siRNA polyplex stability, pharmacokinetics, and potency in vivo. Journal of Controlled Release 2017, 255, 12–26.

11. Boussif, O.; Lezoualc’h, F.; Zanta, M. A.; Mergny, M. D.; Scherman, D.; Demeneix, B.; Behr, J. P., A versatile vector for gene and oligonucleotide transfer into cells in culture and in vivo: polyethylenimine. Proc Natl Acad Sci U S A 1995, 92 (16), 7297–301.

12. Vermeulen, L. M. P.; De Smedt, S. C.; Remaut, K.; Braeckmans, K., The proton sponge hypothesis: Fable or fact? Eur J Pharm Biopharm 2018, 129, 184–190.

13. Winkeljann, B.; Keul, D. C.; Merkel, O. M., Engineering poly- and micelleplexes for nucleic acid delivery - A reflection on their endosomal escape. J Control Release 2022, 353, 518–534.

14. Monnery, B. D., Polycation-Mediated Transfection: Mechanisms of Internalization and Intracellular Trafficking. Biomacromolecules 2021, 22 (10), 4060–4083.

15. Rehman, Z. u.; Hoekstra, D.; Zuhorn, I. S., Mechanism of Polyplex- and Lipoplex-Mediated Delivery of Nucleic Acids: Real-Time Visualization of Transient Membrane Destabilization without Endosomal Lysis. ACS Nano 2013, 7 (5), 3767–3777.

16. Bates, S. M.; Munson, M. J.; Trovisco, V.; Pereira, S.; Miller, S. R.; Sabirsh, A.; Betts, C. J.; Blenke, E. O.; Gay, N. J., The kinetics of endosomal disruption reveal differences in lipid nanoparticle induced cellular toxicity. Journal of Controlled Release 2025, 386.

17. Vaidyanathan, S.; Orr, B. G.; Banaszak Holl, M. M., Role of Cell Membrane–Vector Interactions in Successful Gene Delivery. Accounts of Chemical Research 2016, 49 (8), 1486–1493.

18. Degors, I. M. S.; Wang, C.; Rehman, Z. U.; Zuhorn, I. S., Carriers Break Barriers in Drug Delivery: Endocytosis and Endosomal Escape of Gene Delivery Vectors. Accounts of Chemical Research 2019, 52 (7), 1750–1760.

19. Benjaminsen, R. V.; Mattebjerg, M. A.; Henriksen, J. R.; Moghimi, S. M.; Andresen, T. L., The possible “proton sponge “ effect of polyethylenimine (PEI) does not include change in lysosomal pH. Mol Ther 2013, 21 (1), 149–57.

20. Bruininks, B. M.; Souza, P. C.; Ingolfsson, H.; Marrink, S. J., A molecular view on the escape of lipoplexed DNA from the endosome. Elife 2020, 9.

21. Johansson, J. M.; Du Rietz, H.; Hedlund, H.; Eriksson, H. C.; Oude Blenke, E.; Pote, A.; Harun, S.; Nordenfelt, P.; Lindfors, L.; Wittrup, A., Cellular and biophysical barriers to lipid nanoparticle mediated delivery of RNA to the cytosol. 2024.

22. Spadea, A.; Jackman, M.; Cui, L.; Pereira, S.; Lawrence, M. J.; Campbell, R. A.; Ashford, M., Nucleic Acid-Loaded Lipid Nanoparticle Interactions with Model Endosomal Membranes. ACS Applied Materials & Interfaces 2022, 14 (26), 30371–30384.

23. Omo-Lamai, S.; Wang, Y.; Patel, M. N.; Milosavljevic, A.; Zuschlag, D.; Poddar, S.; Wu, J.; Wang, L.; Dong, F.; Espy, C.; Majumder, A.; Essien, E.-O.; Shen, M.; Channer, B.; Papp, T. E.; Tobin, M.; Maheshwari, R.; Jeong, S.; Patel, S.; Shah, A.; Murali, S.; Chase, L. S.; Zamora, M. E.; Arral, M. L.; Marcos-Contreras, O. A.; Myerson, J. W.; Hunter, C. A.; Discher, D.; Gaskill, P. J.; Tsourkas, A.; Muzykantov, V. R.; Brodsky, I.; Shin, S.; Whitehead, K. A.; Parhiz, H.; Katzen, J.; Miner, J. J.; Trauner, D.; Brenner, J. S., Limiting endosomal damage sensing reduces inflammation triggered by lipid nanoparticle endosomal escape. Nature Nanotechnology 2025.

24. Kristen, A. V.; Ajroud-Driss, S.; Conceicao, I.; Gorevic, P.; Kyriakides, T.; Obici, L., Patisiran, an RNAi therapeutic for the treatment of hereditary transthyretin-mediated amyloidosis. Neurodegener Dis Manag 2019, 9 (1), 5–23.

25. Gilleron, J.; Querbes, W.; Zeigerer, A.; Borodovsky, A.; Marsico, G.; Schubert, U.; Manygoats, K.; Seifert, S.; Andree, C.; Stöter, M.; Epstein-Barash, H.; Zhang, L.; Koteliansky, V.; Fitzgerald, K.; Fava, E.; Bickle, M.; Kalaidzidis, Y.; Akinc, A.; Maier, M.; Zerial, M., Image-based analysis of lipid nanoparticle–mediated siRNA delivery, intracellular trafficking and endosomal escape. Nature Biotechnology 2013, 31 (7), 638–646.

26. Sabnis, S.; Kumarasinghe, E. S.; Salerno, T.; Mihai, C.; Ketova, T.; Senn, J. J.; Lynn, A.; Bulychev, A.; McFadyen, I.; Chan, J.; Almarsson, Ö.; Stanton, M. G.; Benenato, K. E., A Novel Amino Lipid Series for mRNA Delivery: Improved Endosomal Escape and Sustained Pharmacology and Safety in Non-human Primates. Molecular Therapy 2018, 26 (6), 1509–1519.

27. Angelescu, D. G., Molecular modeling of the carbohydrate corona formation on a polyvinyl chloride nanoparticle and its impact on the adhesion to lipid bilayers. J Chem Phys 2024, 160 (14).

28. Grasso, G.; Deriu, M. A.; Patrulea, V.; Borchard, G.; Moller, M.; Danani, A., Free energy landscape of siRNA-polycation complexation: Elucidating the effect of molecular geometry, polymer flexibility, and charge neutralization. PLoS One 2017, 12 (10), e0186816.

29. Zhao, X.; Xu, Q.; Wang, Q.; Liang, X.; Wang, J.; Jin, H.; Man, Y.; Guo, D.; Gao, F.; Tang, X., Induced Self-Assembly of Vitamin E-Spermine/siRNA Nanocomplexes via Spermine/Helix Groove-Specific Interaction for Efficient siRNA Delivery and Antitumor Therapy. Adv Healthc Mater 2024, e2303186.

30. Souza, P. C. T.; Alessandri, R.; Barnoud, J.; Thallmair, S.; Faustino, I.; Grunewald, F.; Patmanidis, I.; Abdizadeh, H.; Bruininks, B. M. H.; Wassenaar, T. A.; Kroon, P. C.; Melcr, J.; Nieto, V.; Corradi, V.; Khan, H. M.; Domanski, J.; Javanainen, M.; Martinez-Seara, H.; Reuter, N.; Best, R. B.; Vattulainen, I.; Monticelli, L.; Periole, X.; Tieleman, D. P.; de Vries, A. H.; Marrink, S. J., Martini 3: a general purpose force field for coarse-grained molecular dynamics. Nat Methods 2021, 18 (4), 382–388.

31. Steinegger, K. M.; Allmendinger, L.; Sturm, S.; Sieber-Schafer, F.; Kromer, A. P. E.; Muller-Caspary, K.; Winkeljann, B.; Merkel, O. M., Molecular Dynamics Simulations Elucidate the Molecular Organization of Poly(beta-amino ester) Based Polyplexes for siRNA Delivery. Nano Lett 2024.

32. Paloncyova, M.; Srejber, M.; Cechova, P.; Kuhrova, P.; Zaoral, F.; Otyepka, M., Atomistic Insights into Organization of RNA-Loaded Lipid Nanoparticles. J Phys Chem B 2023, 127 (5), 1158–1166.

33. Zheng, M.; Pavan, G. M.; Neeb, M.; Schaper, A. K.; Danani, A.; Klebe, G.; Merkel, O. M.; Kissel, T., Targeting the blind spot of polycationic nanocarrier-based siRNA delivery. ACS Nano 2012, 6 (11), 9447–54.

34. Jin, Y.; Adams, F.; Möller, J.; Isert, L.; Zimmermann, C. M.; Keul, D.; Merkel, O. M., Synthesis and Application of Low Molecular Weight PEI-Based Copolymers for siRNA Delivery with Smart Polymer Blends. Macromolecular Bioscience 2022, 23 (2).

35. Kromer, A. P. E.; Sieber-Schafer, F.; Farfan Benito, J.; Merkel, O. M., Design of Experiments Grants Mechanistic Insights into the Synthesis of Spermine-Containing PBAE Copolymers. ACS Appl Mater Interfaces 2024.

36. Kjølbye, L. R.; Valério, M.; Paloncýová, M.; Borges-Araújo, L.; Pestana-Nobles, R.; Grünewald, F.; M. H. Bruininks, B.; Araya-Osorio, R.; Šrejber, M.; Mera-Adasme, R.; Monticelli, L.; J. Marrink, S.; Otyepka, M.; Wu, S.; C.T. Souza, P., Martini 3 building blocks for Lipid Nanoparticle design. ChemRxiv 2025.

37. Müller, J. T.; Kromer, A. P. E.; Ezaddoustdar, A.; Alexopoulos, I.; Steinegger, K. M.; Porras-Gonzalez, D. L.; Berninghausen, O.; Beckmann, R.; Braubach, P.; Burgstaller, G.; Wygrecka, M.; Merkel, O. M., Nebulization of RNA-Loaded Micelle-Embedded Polyplexes as a Potential Treatment of Idiopathic Pulmonary Fibrosis. ACS Applied Materials & Interfaces 2025, 17 (8), 11861–11872.

38. Wilson, S. C.; Baryza, J. L.; Reynolds, A. J.; Bowman, K.; Keegan, M. E.; Standley, S. M.; Gardner, N. P.; Parmar, P.; Agir, V. O.; Yadav, S.; Zunic, A.; Vargeese, C.; Lee, C. C.; Rajan, S., Real Time Measurement of PEG Shedding from Lipid Nanoparticles in Serum via NMR Spectroscopy. Molecular Pharmaceutics 2015, 12 (2), 386–392.

39. Alabi, C. A.; Sahay, G.; Langer, R.; Anderson, D. G., Development of siRNA-probes for studying intracellular trafficking of siRNA nanoparticles. Integrative Biology 2013, 5 (1), 224–230.

40. Wittrup, A.; Ai, A.; Liu, X.; Hamar, P.; Trifonova, R.; Charisse, K.; Manoharan, M.; Kirchhausen, T.; Lieberman, J., Visualizing lipid-formulated siRNA release from endosomes and target gene knockdown. Nat Biotechnol 2015, 33 (8), 870–6.

41. Du Rietz, H.; Hedlund, H.; Wilhelmson, S.; Nordenfelt, P.; Wittrup, A., Imaging small molecule-induced endosomal escape of siRNA. Nature Communications 2020, 11 (1).

42. Boya, P.; Kroemer, G., Lysosomal membrane permeabilization in cell death. Oncogene 2008, 27 (50), 6434–6451.

43. Rui, Y.; Wilson, D. R.; Tzeng, S. Y.; Yamagata, H. M.; Sudhakar, D.; Conge, M.; Berlinicke, C. A.; Zack, D. J.; Tuesca, A.; Green, J. J., High-throughput and high-content bioassay enables tuning of polyester nanoparticles for cellular uptake, endosomal escape, and systemic in vivo delivery of mRNA. Sci Adv 2022, 8 (1), eabk2855.

44. Sieber-Schäfer, F.; Jiang, M.; Kromer, A.; Nguyen, A.; Molbay, M.; Pinto Carneiro, S.; Jürgens, D.; Burgstaller, G.; Popper, B.; Winkeljann, B.; Merkel, O. M., Machine Learning-Enabled Polymer Discovery for Enhanced Pulmonary siRNA Delivery. Advanced Functional Materials 2025.

45. Sæbø, I.; Bjørås, M.; Franzyk, H.; Helgesen, E.; Booth, J., Optimization of the Hemolysis Assay for the Assessment of Cytotoxicity. International Journal of Molecular Sciences 2023, 24 (3).

46. Khang, D.; Lee, Y. K.; Choi, E.-J.; Webster, T. J.; Kim, S.-H., Effect of the protein corona on nanoparticles for modulating cytotoxicity and immunotoxicity. International Journal of Nanomedicine 2014.

47. Darwich, Z.; Klymchenko, A. S.; Dujardin, D.; Mély, Y., Imaging lipid order changes in endosome membranes of live cells by using a Nile Red-based membrane probe. RSC Adv. 2014, 4 (17), 8481–8488.

48. Levental, I.; Levental, K. R.; Heberle, F. A., Lipid Rafts: Controversies Resolved, Mysteries Remain. Trends Cell Biol 2020, 30 (5), 341–353.

49. Hall, A.; Róg, T.; Karttunen, M.; Vattulainen, I., Role of Glycolipids in Lipid Rafts: A View through Atomistic Molecular Dynamics Simulations with Galactosylceramide. The Journal of Physical Chemistry B 2010, 114 (23), 7797–7807.

50. Pogozheva, I. D.; Armstrong, G. A.; Kong, L.; Hartnagel, T. J.; Carpino, C. A.; Gee, S. E.; Picarello, D. M.; Rubin, A. S.; Lee, J.; Park, S.; Lomize, A. L.; Im, W., Comparative Molecular Dynamics Simulation Studies of Realistic Eukaryotic, Prokaryotic, and Archaeal Membranes. J Chem Inf Model 2022, 62 (4), 1036–1051.

51. Gallala, H. D.; Sandhoff, K., Biological Function of the Cellular Lipid BMP—BMP as a Key Activator for Cholesterol Sorting and Membrane Digestion. Neurochemical Research 2010, 36 (9), 1594–1600.

52. Cechova, P.; Paloncyova, M.; Srejber, M.; Otyepka, M., Mechanistic insights into interactions between ionizable lipid nanodroplets and biomembranes. J Biomol Struct Dyn 2024, 1–11.

53. Erazo-Oliveras, A.; Najjar, K.; Truong, D.; Wang, T. Y.; Brock, D. J.; Prater, A. R.; Pellois, J. P., The Late Endosome and Its Lipid BMP Act as Gateways for Efficient Cytosolic Access of the Delivery Agent dfTAT and Its Macromolecular Cargos. Cell Chem Biol 2016, 23 (5), 598–607.

54. Kwolek, U.; Jamróz, D.; Janiczek, M.; Nowakowska, M.; Wydro, P.; Kepczynski, M., Interactions of Polyethylenimines with Zwitterionic and Anionic Lipid Membranes. Langmuir 2016, 32 (19), 5004–5018.

55. Wang, C.; Zhao, T.; Li, Y.; Huang, G.; White, M. A.; Gao, J., Investigation of endosome and lysosome biology by ultra pH-sensitive nanoprobes. Advanced Drug Delivery Reviews 2017, 113, 87–96.

56. Omo-Lamai, S.; Wang, Y.; Patel, M. N.; Essien, E.-O.; Shen, M.; Majumdar, A.; Espy, C.; Wu, J.; Channer, B.; Tobin, M.; Murali, S.; Papp, T. E.; Maheshwari, R.; Wang, L.; Chase, L. S.; Zamora, M. E.; Arral, M. L.; Marcos-Contreras, O. A.; Myerson, J. W.; Hunter, C. A.; Tsourkas, A.; Muzykantov, V.; Brodsky, I.; Shin, S.; Whitehead, K. A.; Gaskill, P.; Discher, D.; Parhiz, H.; Brenner, J. S., Lipid Nanoparticle-Associated Inflammation is Triggered by Sensing of Endosomal Damage: Engineering Endosomal Escape Without Side Effects. 2024.

57. Cullis, P. R.; Hope, M. J., Effects of fusogenic agent on membrane structure of erythrocyte ghosts and the mechanism of membrane fusion. Nature 1978, 271 (5646), 672–674.

58. Brosio, G.; Rossi, G.; Bochicchio, D., Nanoparticle-induced biomembrane fusion: unraveling the effect of core size on stalk formation. Nanoscale Advances 2023, 5 (18), 4675–4680.

59. Sabin, J.; Alatorre-Meda, M.; Minones, J., Jr.; Dominguez-Arca, V.; Prieto, G., New insights on the mechanism of polyethylenimine transfection and their implications on gene therapy and DNA vaccines. Colloids Surf B Biointerfaces 2022, 210, 112219.

60. Sarett, S. M.; Werfel, T. A.; Chandra, I.; Jackson, M. A.; Kavanaugh, T. E.; Hattaway, M. E.; Giorgio, T. D.; Duvall, C. L., Hydrophobic interactions between polymeric carrier and palmitic acid-conjugated siRNA improve PEGylated polyplex stability and enhance in vivo pharmacokinetics and tumor gene silencing. Biomaterials 2016, 97, 122–132.

61. Grigsby, C. L.; Leong, K. W., Balancing protection and release of DNA: tools to address a bottleneck of non-viral gene delivery. Journal of The Royal Society Interface 2009, 7 (suppl_1).

62. Habeeb, A. F. S. A., Determination of free amino groups in proteins by trinitrobenzenesulfonic acid. Analytical Biochemistry 1966, 14 (3), 328–336.

63. Petersen, H.; Fechner, P. M.; Martin, A. L.; Kunath, K.; Stolnik, S.; Roberts, C. J.; Fischer, D.; Davies, M. C.; Kissel, T., Polyethylenimine-graft-Poly(ethylene glycol) Copolymers:□ Influence of Copolymer Block Structure on DNA Complexation and Biological Activities as Gene Delivery System. Bioconjugate Chemistry 2002, 13 (4), 845–854.

64. Van Der Spoel, D.; Lindahl, E.; Hess, B.; Groenhof, G.; Mark, A. E.; Berendsen, H. J., GROMACS: fast, flexible, and free. J Comput Chem 2005, 26 (16), 1701–18.

65. Essmann, U.; Perera, L.; Berkowitz, M. L.; Darden, T.; Lee, H.; Pedersen, L. G., A smooth particle mesh Ewald method. The Journal of Chemical Physics 1995, 103 (19), 8577–8593.

66. Parrinello, M.; Rahman, A., Polymorphic transitions in single crystals: A new molecular dynamics method. Journal of Applied Physics 1981, 52 (12), 7182–7190.

67. Binder, J.; Winkeljann, J.; Steinegger, K.; Trnovec, L.; Orekhova, D.; Zahringer, J.; Horner, A.; Fell, V.; Tinnefeld, P.; Winkeljann, B.; Friess, W.; Merkel, O. M., Closing the Gap between Experiment and Simulation: A Holistic Study on the Complexation of Small Interfering RNAs with Polyethylenimine. Mol Pharm 2024, 21 (5), 2163–2175.

68. Jorgensen, W. L.; Tirado-Rives, J., Potential energy functions for atomic-level simulations of water and organic and biomolecular systems. Proc Natl Acad Sci U S A 2005, 102 (19), 6665–70.

69. Dodda, L. S.; Vilseck, J. Z.; Tirado-Rives, J.; Jorgensen, W. L., 1.14*CM1A-LBCC: Localized Bond-Charge Corrected CM1A Charges for Condensed-Phase Simulations. J Phys Chem B 2017, 121 (15), 3864–3870.

70. Dodda, L. S.; Cabeza de Vaca, I.; Tirado-Rives, J.; Jorgensen, W. L., LigParGen web server: an automatic OPLS-AA parameter generator for organic ligands. Nucleic Acids Res 2017, 45 (W1), W331–W336.

71. Barnoud, J. https://github.com/jbarnoud/cgbuilder.

72. Grava, M.; Ibrahim, M.; Sudarsan, A.; Pusterla, J.; Philipp, J.; Radler, J. O.; Schwierz, N.; Schneck, E., Combining molecular dynamics simulations and x-ray scattering techniques for the accurate treatment of protonation degree and packing of ionizable lipids in monolayers. J Chem Phys 2023, 159 (15).

73. Sezgin, E.; Levental, I.; Mayor, S.; Eggeling, C., The mystery of membrane organization: composition, regulation and roles of lipid rafts. Nat Rev Mol Cell Biol 2017, 18 (6), 361–374.

74. Escriba, P. V.; Busquets, X.; Inokuchi, J.; Balogh, G.; Torok, Z.; Horvath, I.; Harwood, J. L.; Vigh, L., Membrane lipid therapy: Modulation of the cell membrane composition and structure as a molecular base for drug discovery and new disease treatment. Prog Lipid Res 2015, 59, 38–53.

75. Yang, Y.; Lee, M.; Fairn, G. D., Phospholipid subcellular localization and dynamics. J Biol Chem 2018, 293 (17), 6230–6240.

76. Zaborowska, M.; Dziubak, D.; Matyszewska, D.; Sek, S.; Bilewicz, R., Designing a Useful Lipid Raft Model Membrane for Electrochemical and Surface Analytical Studies. Molecules 2021, 26 (18).

77. Krause, M. R.; Regen, S. L., The structural role of cholesterol in cell membranes: from condensed bilayers to lipid rafts. Acc Chem Res 2014, 47 (12), 3512–21.

78. Wassenaar, T. A.; Ingolfsson, H. I.; Bockmann, R. A.; Tieleman, D. P.; Marrink, S. J., Computational Lipidomics with insane: A Versatile Tool for Generating Custom Membranes for Molecular Simulations. J Chem Theory Comput 2015, 11 (5), 2144–55.

79. Lopez, C. A.; Sovova, Z.; van Eerden, F. J.; de Vries, A. H.; Marrink, S. J., Martini Force Field Parameters for Glycolipids. J Chem Theory Comput 2013, 9 (3), 1694–708.

80. Borges-Araujo, L.; Souza, P. C. T.; Fernandes, F.; Melo, M. N., Improved Parameterization of Phosphatidylinositide Lipid Headgroups for the Martini 3 Coarse-Grain Force Field. J Chem Theory Comput 2022, 18 (1), 357–373.

81. Grunewald, F.; Punt, M. H.; Jefferys, E. E.; Vainikka, P. A.; Konig, M.; Virtanen, V.; Meyer, T. A.; Pezeshkian, W.; Gormley, A. J.; Karonen, M.; Sansom, M. S. P.; Souza, P. C. T.; Marrink, S. J., Martini 3 Coarse-Grained Force Field for Carbohydrates. J Chem Theory Comput 2022, 18 (12), 7555–7569.

82. Calandrini, V.; Pellegrini, E.; Calligari, P.; Hinsen, K.; Kneller, G. R., nMoldyn - Interfacing spectroscopic experiments, molecular dynamics simulations and models for time correlation functions. École thématique de la Société Française de la Neutronique 2011, 12, 201–232.

83. de Buyl, P., tidynamics: A tiny package to compute the dynamics of stochastic and molecular simulations. Journal of Open Source Software 2018, 3 (28).

84. Gowers, R.; Linke, M.; Barnoud, J.; Reddy, T.; Melo, M.; Seyler, S.; Domański, J.; Dotson, D.; Buchoux, S.; Kenney, I.; Beckstein, O., MDAnalysis: A Python Package for the Rapid Analysis of Molecular Dynamics Simulations. In Proceedings of the 15th Python in Science Conference, 2016; pp 98–105.

85. Michaud - Agrawal, N.; Denning, E. J.; Woolf, T. B.; Beckstein, O., MDAnalysis: A toolkit for the analysis of molecular dynamics simulations. Journal of Computational Chemistry 2011, 32 (10), 2319–2327.

86. Torrie, G. M.; Valleau, J. P., Nonphysical sampling distributions in Monte Carlo free-energy estimation: Umbrella sampling. Journal of Computational Physics 1977, 23 (2), 187–199.

87. Fornasier, F.; Souza, L. M. P.; Souza, F. R.; Reynaud, F.; Pimentel, A. S., Lipophilicity of Coarse-Grained Cholesterol Models. J Chem Inf Model 2020, 60 (2), 569–577.

88. Hub, J. S.; de Groot, B. L.; van der Spoel, D., g_wham—A Free Weighted Histogram Analysis Implementation Including Robust Error and Autocorrelation Estimates. Journal of Chemical Theory and Computation 2010, 6 (12), 3713–3720.

89. Kumar, S.; Rosenberg, J. M.; Bouzida, D.; Swendsen, R. H.; Kollman, P. A., THE weighted histogram analysis method for free - energy calculations on biomolecules. I. The method. Journal of Computational Chemistry 2004, 13 (8), 1011–1021.

90. Stelzl, L. S.; Kells, A.; Rosta, E.; Hummer, G., Dynamic Histogram Analysis To Determine Free Energies and Rates from Biased Simulations. Journal of Chemical Theory and Computation 2017, 13 (12), 6328–6342.

91. Jo, S.; Kim, T.; Iyer, V. G.; Im, W., CHARMM-GUI: A web-based graphical user interface for CHARMM. Journal of Computational Chemistry 2008, 29 (11), 1859–1865.

92. Brooks, B. R.; Brooks, C. L.; Mackerell, A. D.; Nilsson, L.; Petrella, R. J.; Roux, B.; Won, Y.; Archontis, G.; Bartels, C.; Boresch, S.; Caflisch, A.; Caves, L.; Cui, Q.; Dinner, A. R.; Feig, M.; Fischer, S.; Gao, J.; Hodoscek, M.; Im, W.; Kuczera, K.; Lazaridis, T.; Ma, J.; Ovchinnikov, V.; Paci, E.; Pastor, R. W.; Post, C. B.; Pu, J. Z.; Schaefer, M.; Tidor, B.; Venable, R. M.; Woodcock, H. L.; Wu, X.; Yang, W.; York, D. M.; Karplus, M., CHARMM: The biomolecular simulation program. Journal of Computational Chemistry 2009, 30 (10), 1545–1614.

93. Lee, J.; Cheng, X.; Swails, J. M.; Yeom, M. S.; Eastman, P. K.; Lemkul, J. A.; Wei, S.; Buckner, J.; Jeong, J. C.; Qi, Y.; Jo, S.; Pande, V. S.; Case, D. A.; Brooks, C. L.; MacKerell, A. D.; Klauda, J. B.; Im, W., CHARMM-GUI Input Generator for NAMD, GROMACS, AMBER, OpenMM, and CHARMM/OpenMM Simulations Using the CHARMM36 Additive Force Field. Journal of Chemical Theory and Computation 2015, 12 (1), 405–413.

94. Wu, E. L.; Cheng, X.; Jo, S.; Rui, H.; Song, K. C.; Dávila-Contreras, E. M.; Qi, Y.; Lee, J.; Monje-Galvan, V.; Venable, R. M.; Klauda, J. B.; Im, W., CHARMM-GUIMembrane Buildertoward realistic biological membrane simulations. Journal of Computational Chemistry 2014, 35 (27), 1997–2004.

95. Jo, S.; Lim, J. B.; Klauda, J. B.; Im, W., CHARMM-GUI Membrane Builder for Mixed Bilayers and Its Application to Yeast Membranes. Biophysical Journal 2009, 97 (1), 50–58.

96. Yuan, A.; Jo, S.; Kim, T.; Im, W., Automated Builder and Database of Protein/Membrane Complexes for Molecular Dynamics Simulations. PLoS ONE 2007, 2 (9).

97. Kim, S.; Lee, J.; Jo, S.; Brooks, C. L.; Lee, H. S.; Im, W., CHARMM-GUI ligand reader and modeler for CHARMM force field generation of small molecules. Journal of Computational Chemistry 2017, 38 (21), 1879–1886.

