## Supplementary Data for "Proton sponge or membrane fusion? – Endosomal escape of siRNA polyplexes illuminated by molecular dynamics simulations"

**Table S1. Composition of model membranes [%]**

Percentage of lipid components by headgroup type in the four different simulated membrane types. Anionic lipids are marked in blue.

| | Early Endosome | | Lipid Raft | | Late Endosome | | Lysosome | |
|---|---|---|---|---|---|---|---|---|
| **% Cholesterol** | | **29.5** | | **45.0** | | **20.1** | | **20.1** |
| Phosphatidylcholines (PC) | POPC | 17.0 | POPC | 27.5 | POPC | 11.0 | POPC | 11.0 |
| | PIPC | 15.1 | | | PIPC | 18.1 | PIPC | 18.1 |
| | PAPC | 5.5 | | | PAPC | 6.0 | PAPC | 6.0 |
| Phosphatidylethanolamines (PE) | PIPE | 4.1 | | | POPE | 11.5 | POPE | 11.5 |
| | PAPE | 6.8 | | | PAPE | 11.5 | PAPE | 11.5 |
| Sphingomyelins (SM) | DPSM | 7.1 | DPSM | 25.5 | DPSM | 2.7 | DPSM | 2.9 |
| | PGSM | 7.1 | | | DXSM | 2.7 | DXSM | 2.9 |
| Phosphatidylserines (PS) | | | | | POPS | 2.0 | POPS | 2.0 |
| Phosphatidylinositols (PI) | PIPI | 7.3 | | | PIPI | 8.0 | PIPI | 8.0 |
| Glycolip-monosialohexosylganglioside (DPG) | DPG1 | 0.2 | DPG1 | 1.0 | DPG1 | 0.2 | | |
| | DPG3 | 0.2 | DPG3 | 1.0 | DPG3 | 0.2 | | |
| Bis(monoacylglycero)phosphate (BMGP) | | | | | BMGP | 6.0 | BMGP | 6.0 |
| **% neg. lipids (sum)** | | **7.7** | | **2.0** | | **16.4** | | **15.9** |

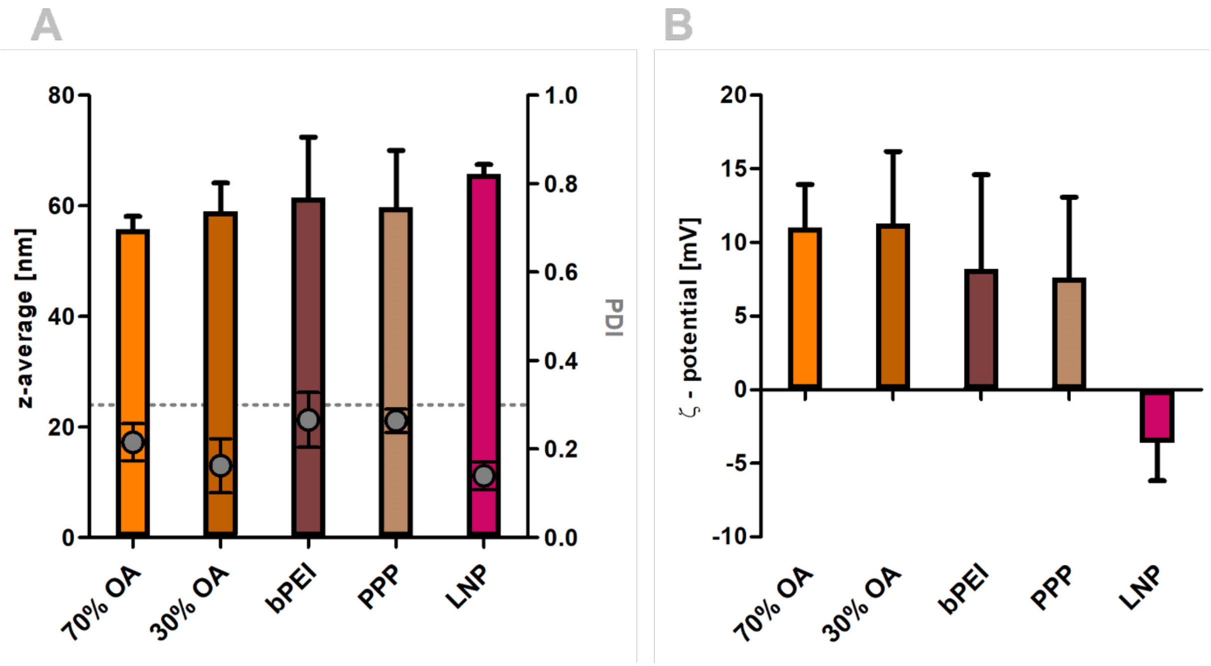


**Figure S1. Size and ζ-potential of nanoparticles A.** Bars: Hydrodynamic diameters shown as z-average in nm, mean ± sd, n = 3; Dots: PDI, mean ± sd, n = 3. **B.** ζ-Potential in mV, mean ± sd, n = 3.

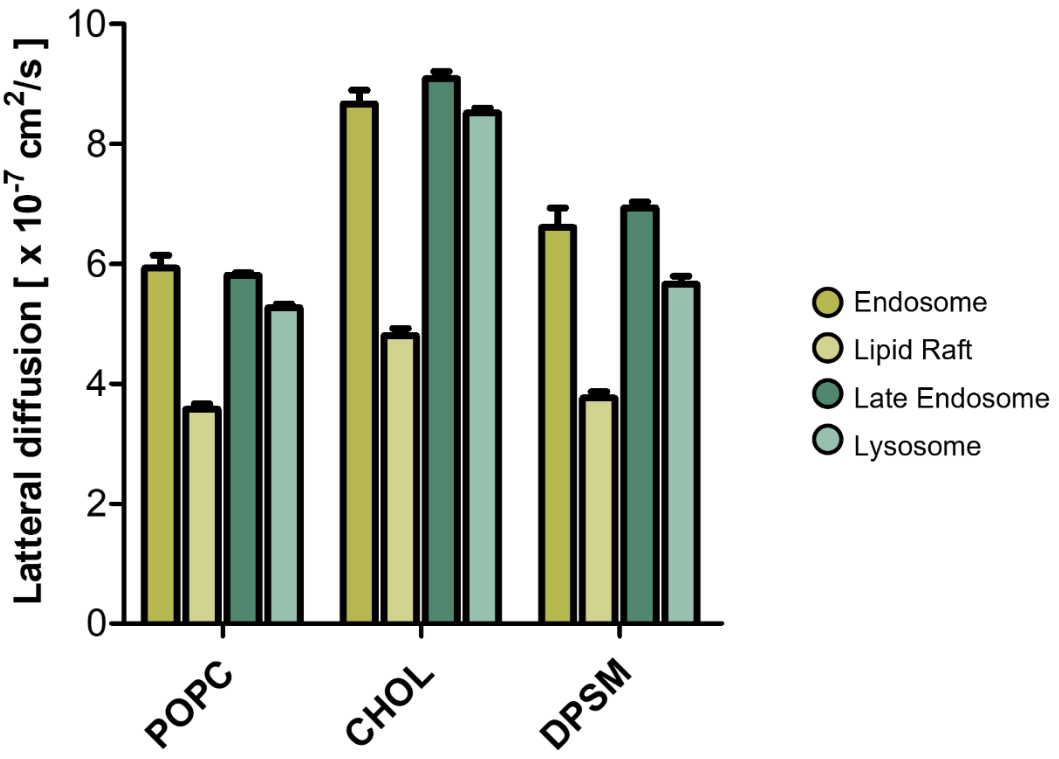


**Figure S2. Lateral diffusion [x 10-7 $cm^2/s$] of membrane lipids in the different CG membrane models.**

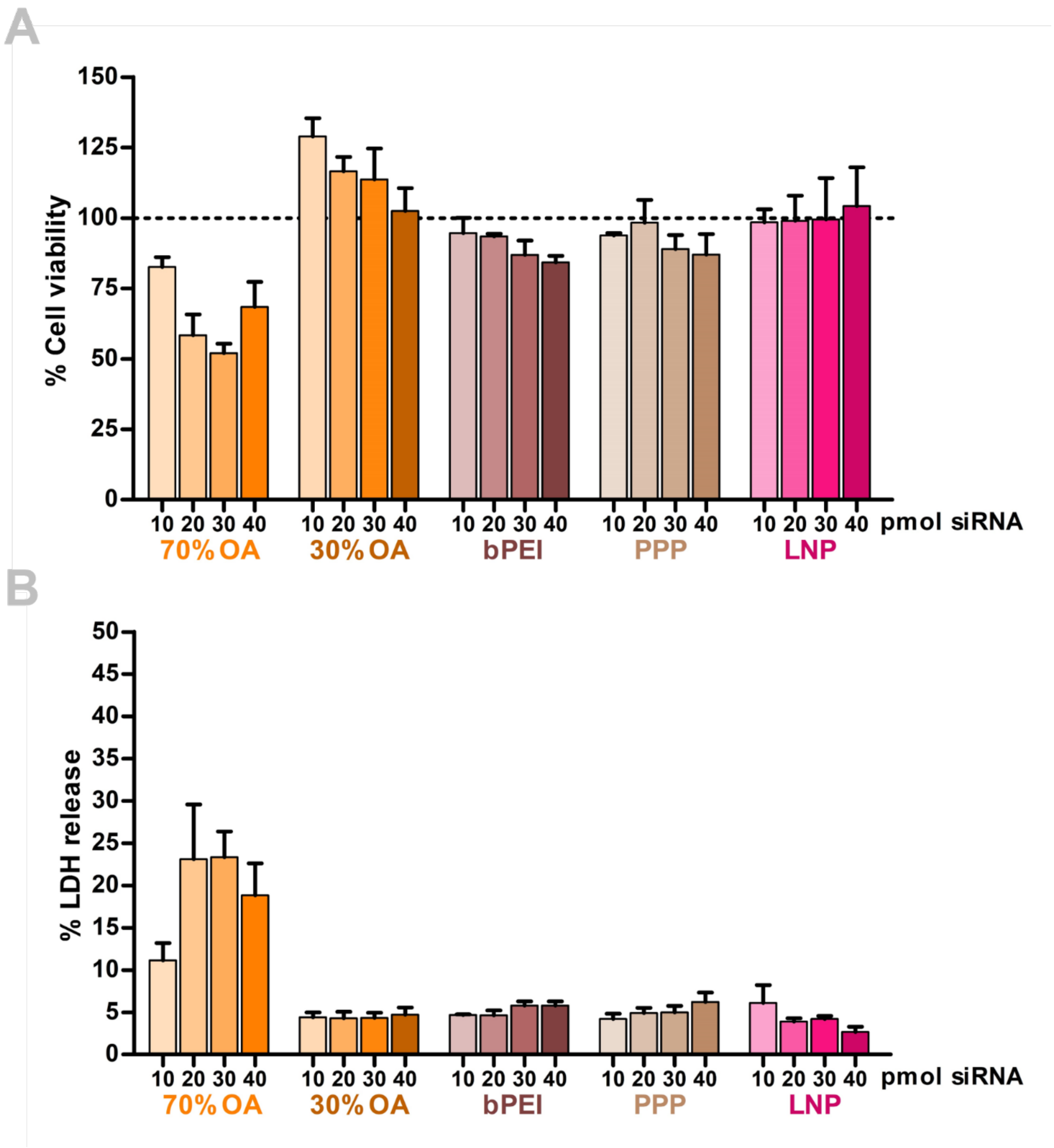


**Figure S3. CCK8 and LDH release assay in HeLa cells. A.** Cell viability [%] calculated from CCK8 assay with 4 different nanoparticle concentrations, mean ± sd, n = 3. **B.** LDH release [%] calculated from the same 4 concentrations of nanoparticles, mean ± sd, n = 3.

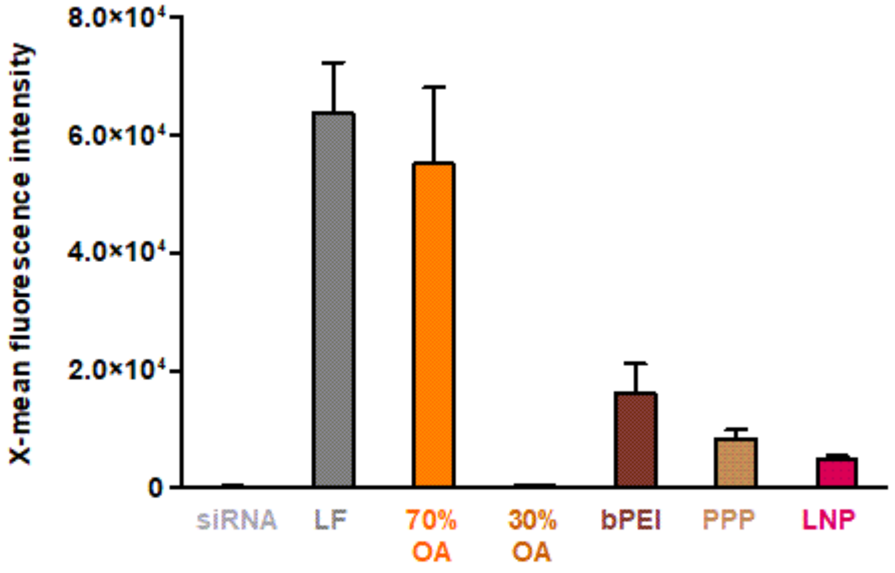


**Figure S4. Cellular uptake in HeLa-eGFP cells, 24 h after transfection with 20 pmol of nanoparticles containing 20% AF647-labelled siGFP, mean ± sd, n = 3.**

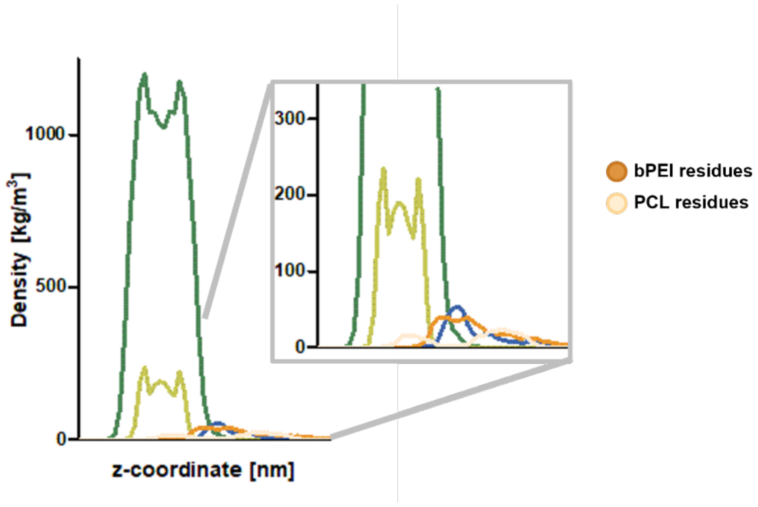


**Figure S5. Density distribution [kg/m$^3$] after interaction of PPP polyplexes and late endosomal membranes.**

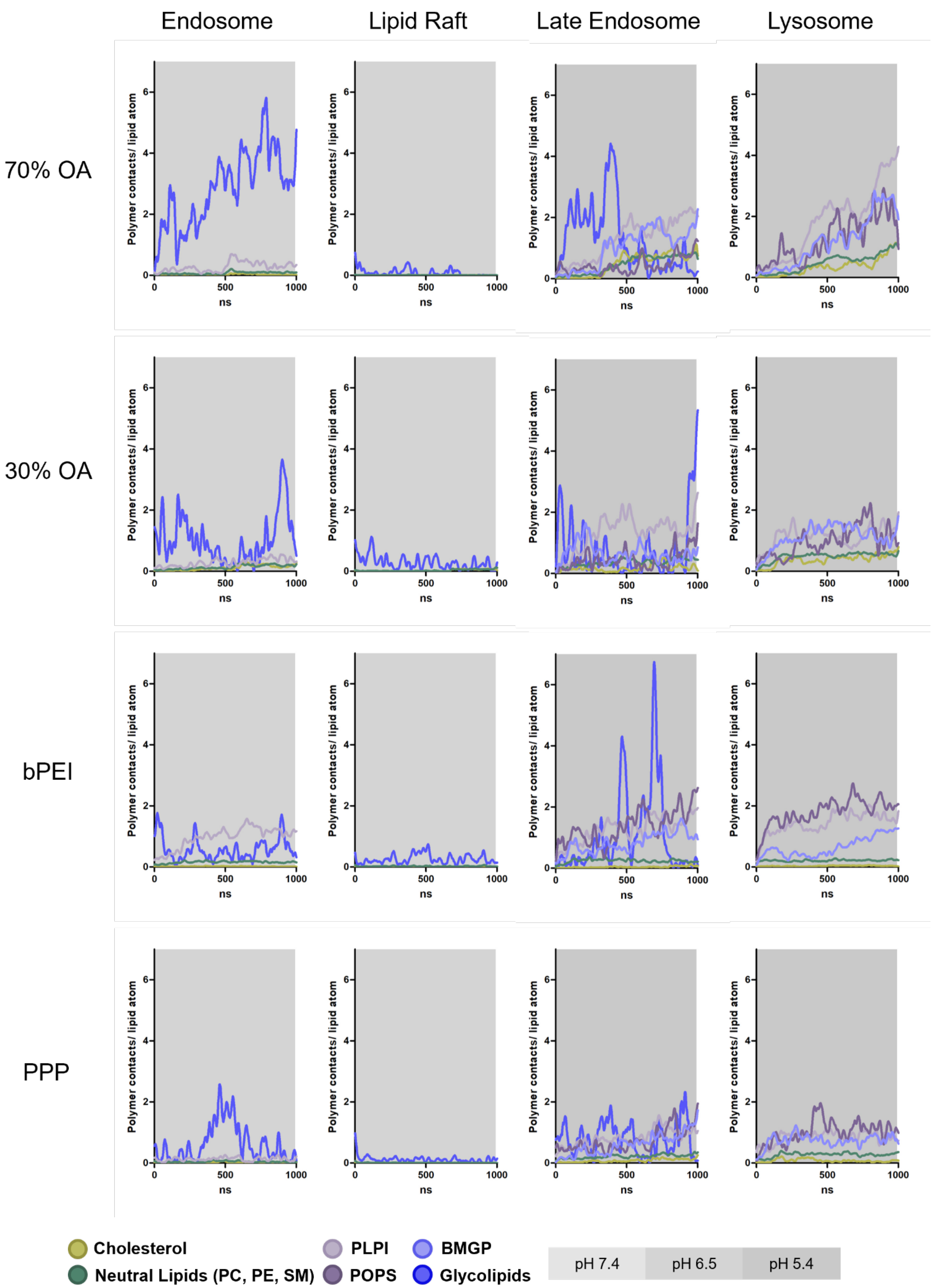


**Figure S6. Interaction of polymer models (in total 9 kDa polymer per simulation box) with planar membranes at AA resolution: Polymer contacts below 0.6 nm per membrane lipid atom over time, n = 2.**

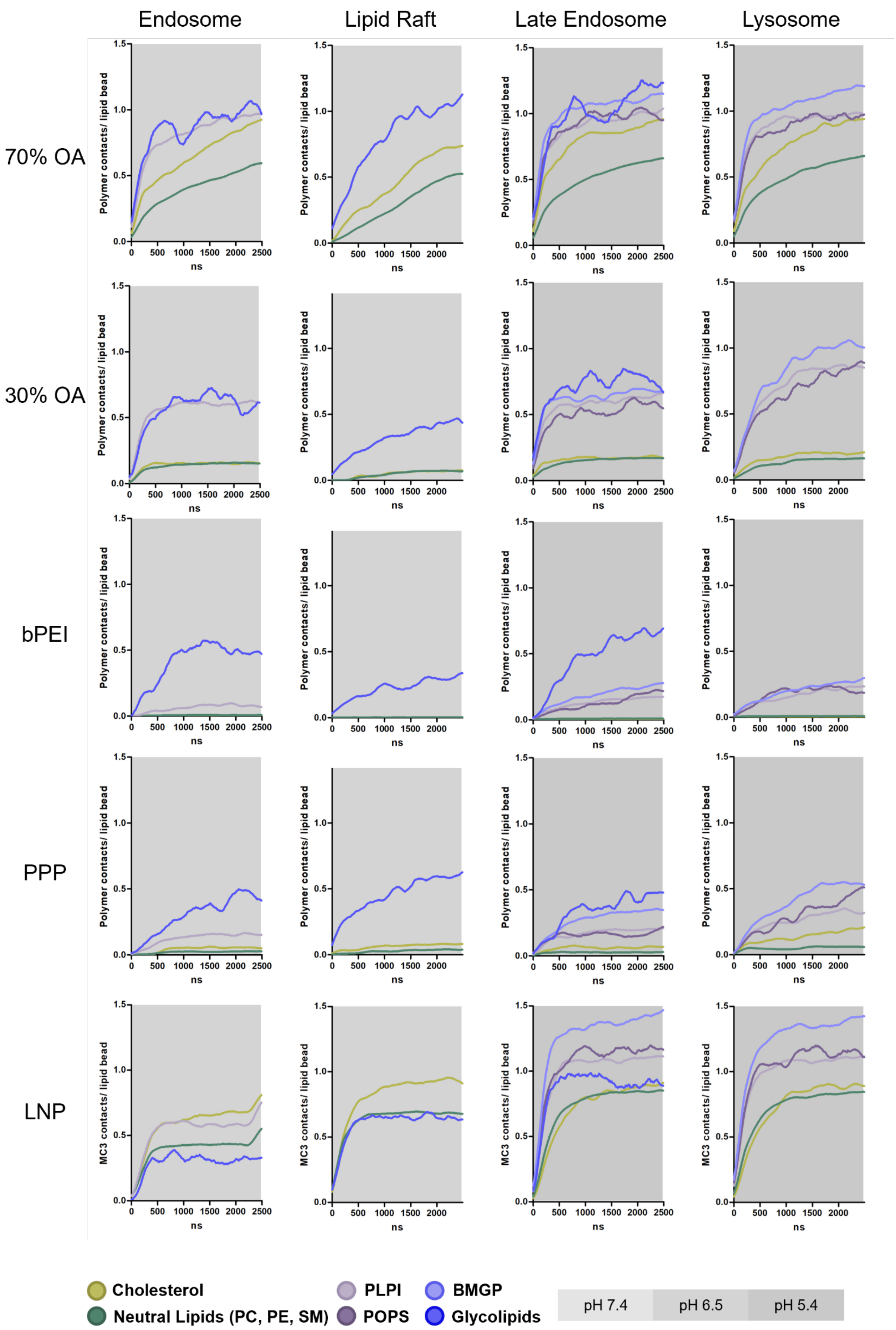


**Figure S7. Interaction of nanoparticles and planar membranes at CG resolution: Polymer or MC3 lipid contacts below 0.6 nm per membrane lipid bead over time, n = 3.**

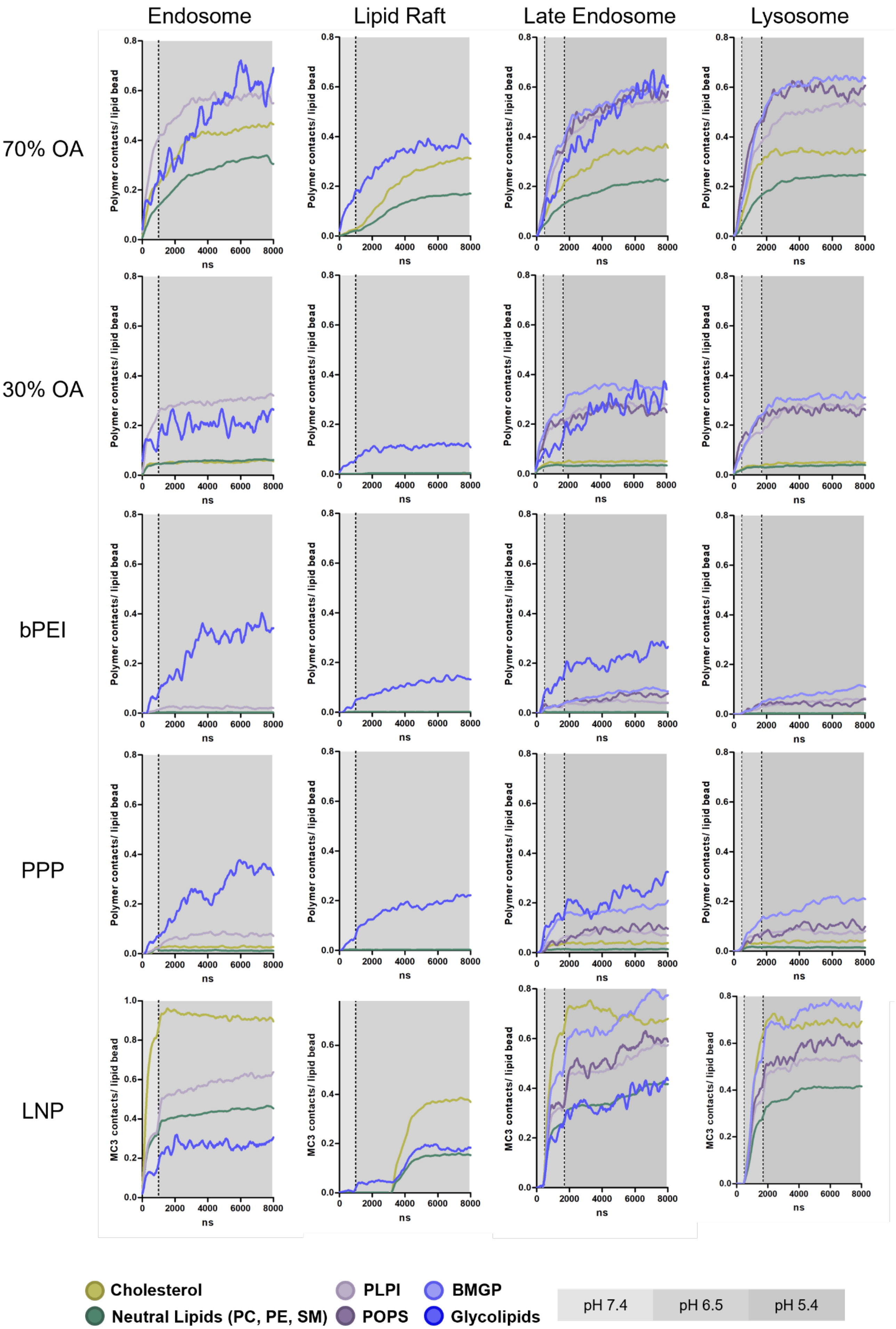


**Figure S8. Interaction of nanoparticles and vesicles in CG resolution: Polymer or MC3 lipid contacts below 0.6 nm per membrane lipid bead over time, n = 2.**

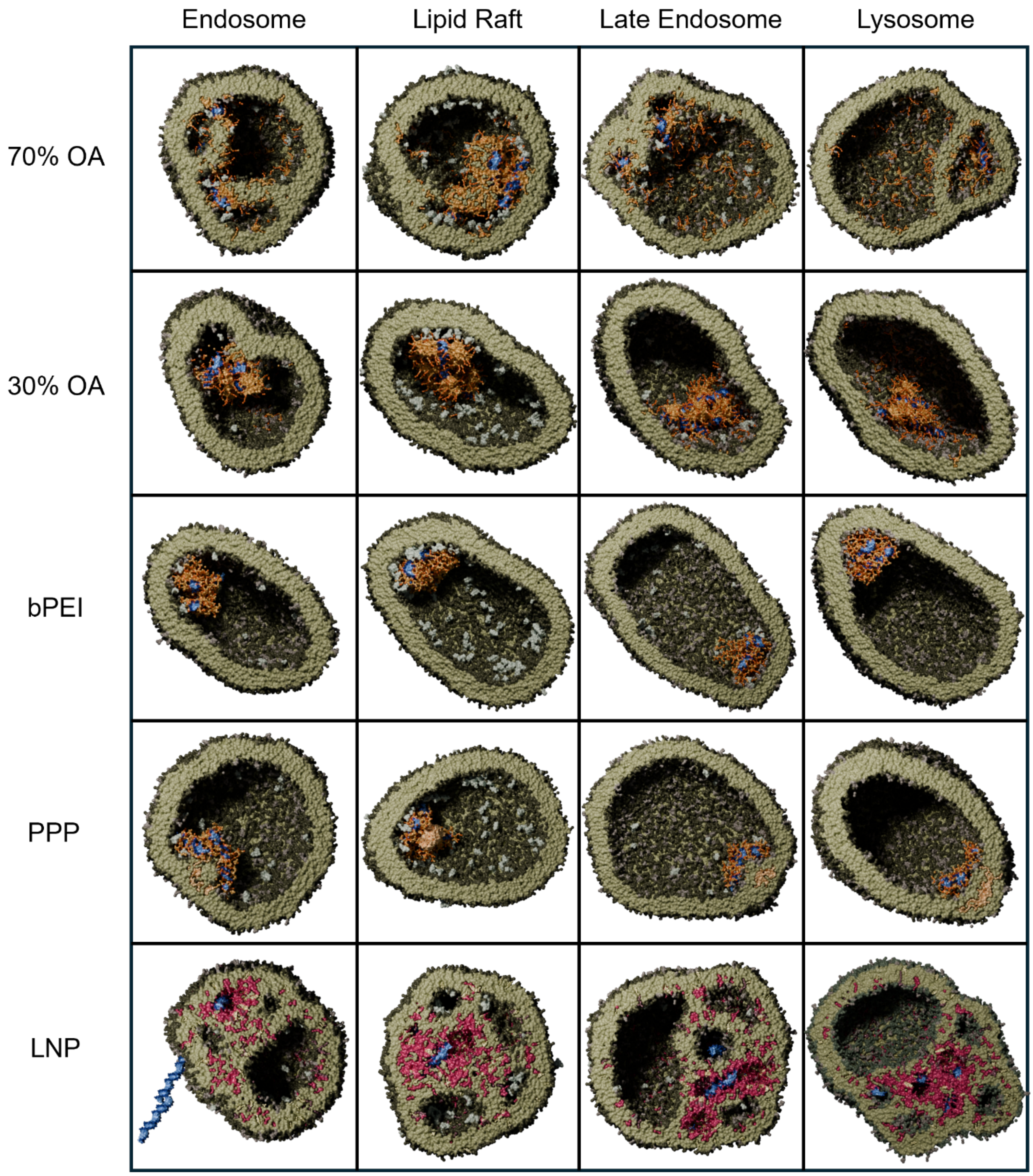


**Figure S9. Matrix of the five nanoparticles interacting with the four different vesicles.**

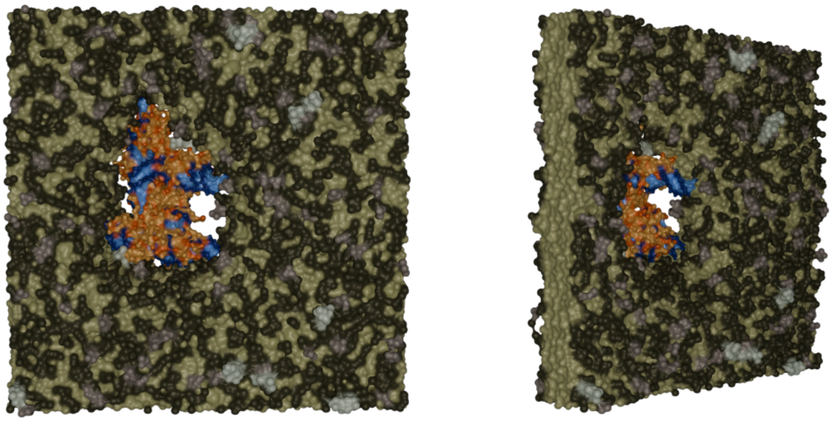

**Figure S10. CG simulation of a bPEI polyplex being forcefully pulled through an endosomal membrane.**

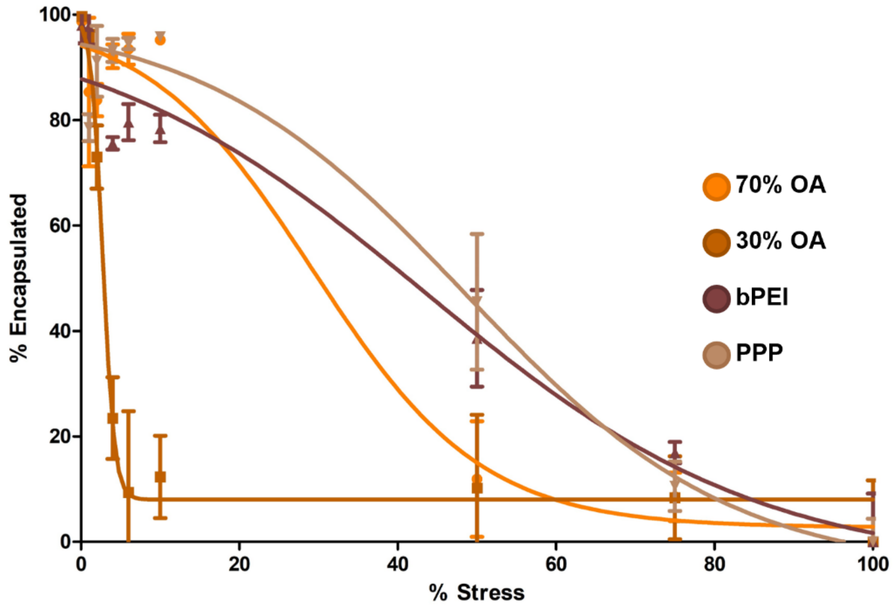


| Polymer | EC50 | $R^2$ |
|---|---|---|
| 70% OA PBAE | 29.4 | 0.963 |
| 30% OA PBAE | 2.75 | 0.953 |
| bPEI | 43.4 | 0.949 |
| PPP | 48.5 | 0.959 |

**Figure S11. Experimental stability data of the polyplex formulations.** % Encapsulated siRNA depending on the concentration (% stress) of the applied heparin/Triton X solution (mean ± sd, n = 3), and sigmoidal fit yielding an EC50 value for each polymer (i.e., the % stress at which 50% of the siRNA is unpacked).
